# Cell-binding IgM in CSF is distinctive of multiple sclerosis and targets the iron transporter SCARA5

**DOI:** 10.1101/2023.09.29.560121

**Authors:** I. Callegari, J. Oechtering, M. Schneider, S. Perriot, A. Mathias, M.M. Voortman, A. Cagol, U. Lanner, M. Diebold, S. Holdermann, V. Kreiner, B. Becher, C. Granziera, A. Junker, R. Du Pasquier, M. Khalil, J. Kuhle, L. Kappos, NSR Sanderson, T. Derfuss

**Author notes:** Correspondence to: Nicholas SR Sanderson, Full address Department of Biomedicine, Hebelstrasse 20, 4056 Basel. These authors contributed equally to this work (NSR Sanderson, T. Derfuss).

## Abstract

Intrathecal IgM production in multiple sclerosis (MS) is associated with a worse disease course. To investigate pathogenic relevance of autoreactive IgM in MS, CSF from two independent cohorts, including MS patients and controls, were screened for antibody binding to induced pluripotent stem cell-derived neurons and astrocytes, and a panel of CNS-related cell lines. IgM binding to a primitive neuro-ectodermal tumour cell line discriminated 10% of MS donors from controls. Transcriptomes of single IgM producing CSF B cells from patients with cell-binding IgM were sequenced and used to produce recombinant monoclonal antibodies for characterisation and antigen identification. We produced 5 cell-binding recombinant IgM antibodies, of which one, cloned from an HLA-DR+ plasma-like B cell, mediated antigen-dependent complement activation. Immunoprecipitation and mass spectrometry, and biochemical and transcriptome analysis of the target cells identified the iron transport scavenger protein SCARA5 as the antigen target of this antibody. Intrathecal injection of a SCARA5 antibody led to an increased T cell infiltration in an EAE model. CSF IgM might contribute to CNS inflammation in MS by binding to cell surface antigens like SCARA5 and activating complement, or by facilitating immune cell migration into the brain.

## Introduction

One of the diagnostic hallmarks of multiple sclerosis (MS) is intrathecal immunoglobulin production, identifiable at CSF examination as oligoclonal IgG bands (OCB). Other than in MS, OCB are found in CNS inflammatory conditions, and when the cause is infectious, antibodies that are specific for the causal pathogen can be identified amongst those constituting the OCB^1,2^.

The prevalent class of intrathecally produced antibodies is IgG^3^, but intrathecally produced immunoglobulin M (IgM) is also present in 20–25% of patients^4^, and their presence has been associated with more active and severe disease^5–8^. IgM sequences from the CSF of patients with MS or neuroborreliosis show a high degree of somatic hypermutation, suggesting an antigen-driven IgM response^9^, rather than a bystander response. In MS, comparison of the immunoglobulin transcriptomes of B cells in CSF and in lesions with peptide sequences of soluble CSF antibodies suggests that all three are clonally related^10,11^. Known mechanisms by which IgM might damage brain tissue are dependent on antigen recognition and binding, implying that if IgM is causally involved in pathogenesis, it must bind antigens in the CNS. By analogy with established pathogenic autoantibodies, we expect such antibodies to bind conformational epitopes on the surface of live cells^12^. A result consistent with this prediction was indeed observed by Lily et al.^13^ who saw that antibodies from serum or CSF from patients with MS had a stronger propensity to bind to human oligodendrocyte and neuronal cell types than samples from healthy patients.

We therefore screened CSF from patients with MS and controls for antibodies of different classes that bind to live, CNS-related cell types. Having discovered an MS-restricted IgM binding pattern, we then set about identifying the target and characterising the pathogenic B cell phenotype.

## Materials and methods

### Methods

Methods are described in detail in the Supplementary Methods section.

### Patient samples

CSF samples from patients undergoing diagnostic lumbar puncture at the University Hospital Basel and from University Hospital of Graz were collected and immediately processed. All procedures were approved by the Ethical Committee of Northwest Switzerland (study protocol 2021-00908) and by the Ethical Committee of the Medical University of Graz, Austria (study protocol 17-046 ex 05/06 and 31-432 ex 18/19). Each sample was collected using a 15 ml falcon tube, centrifuged at 400 g at room temperature for 10 minutes; the CSF supernatant was then aliquoted and frozen at –80°C. For the prospective part of the study, the CSF samples were analysed for cell-line-binding IgM by flow cytometry, immediately after collection, and the CSF cell pellet was resuspended in RPMI supplemented with 40% FCS (R40) and kept on ice for further processing or stored within 2h in liquid nitrogen.

### Cell lines

Cell lines used are described in Supplementary Table 1.

### Statistics

Differences between conditions were tested for deviation from normality, when normal were subjected to analysis of variance, and when significantly deviated from normal, to appropriate non-parametric tests as specified in the figure legends.

## Results

### IgM but not IgG binding to PNET cells discriminates MS patients from controls

First we screened CSF samples from 128 patients with MS and 142 controls for IgM binding to neurons and astrocytes derived from human induced pluripotent stem cells (iPSC) by flow cytometry, blinded to the diagnosis. Overall IgM binding was not obviously different between patients and controls (Fig. 1A), but some samples showed an MS-specific pattern of antibody binding to a subset of cells (Supp Fig. 1). This subset was too small to be easily used for downstream studies, and we therefore screened the same cohort of CSF, plus an additional 44 MS samples and 50 control samples (details in Supplementary Table 2) against a panel of CNS-related tumour cell lines, shown in Supplementary Table 1.

**Figure 1.**
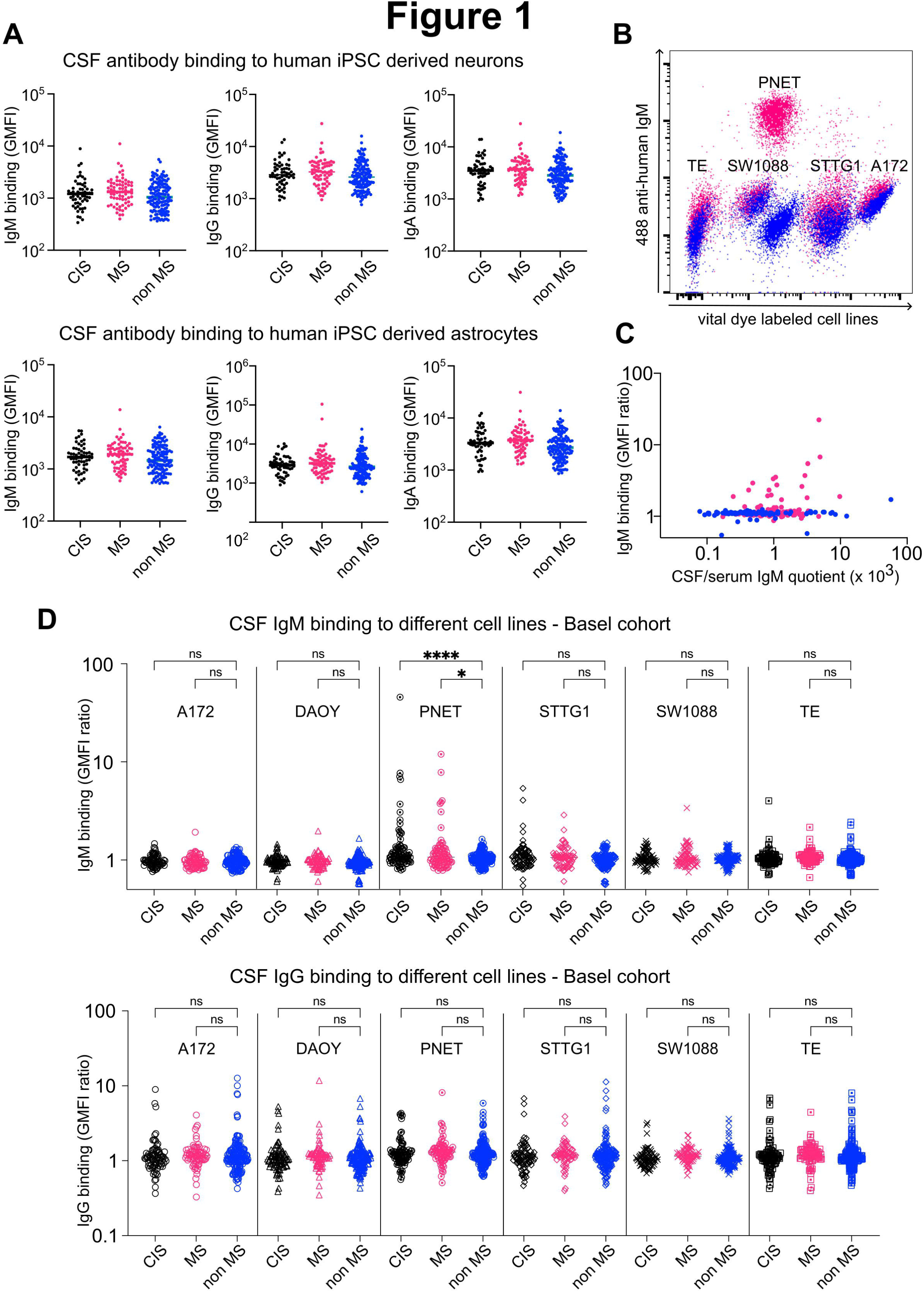
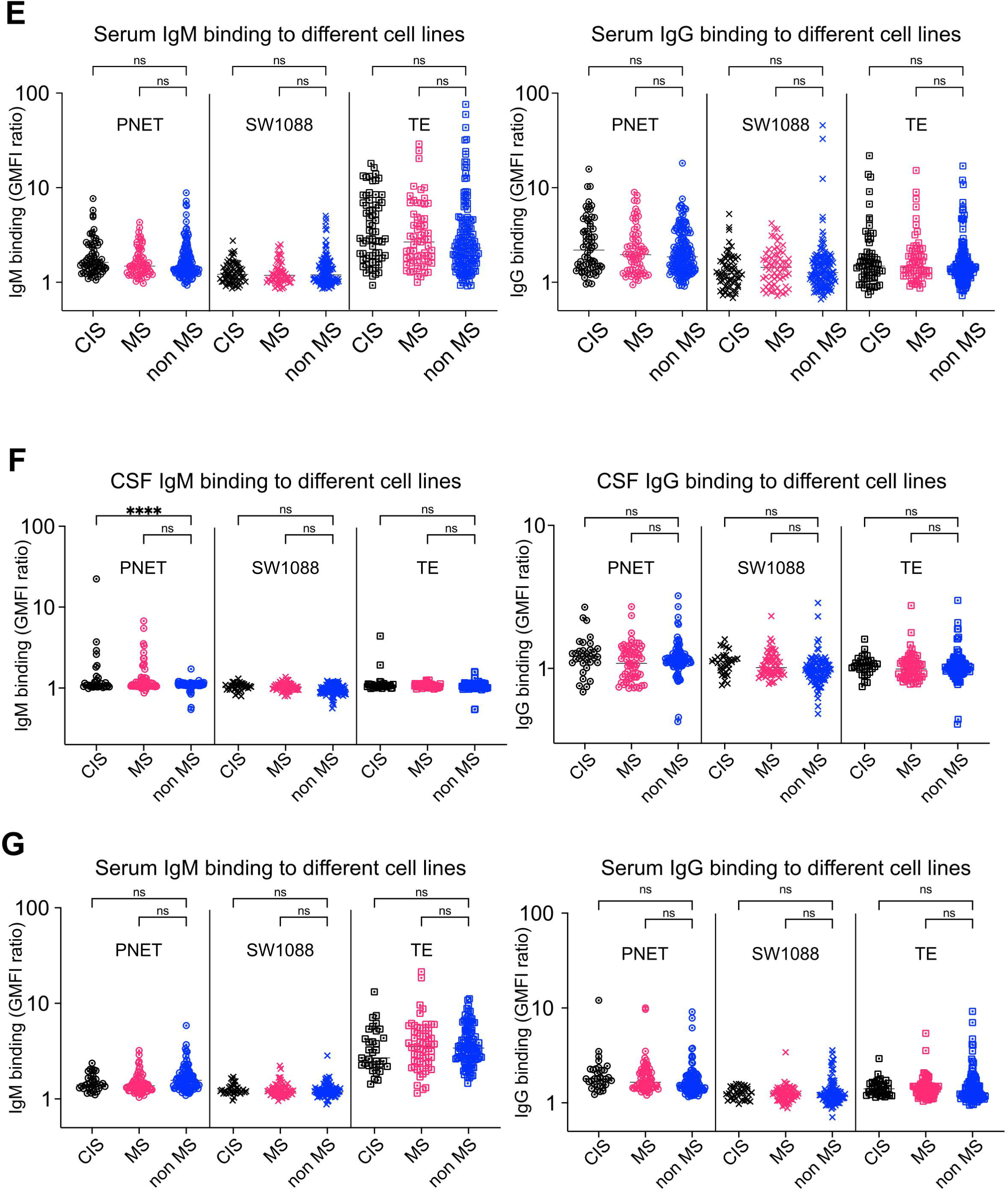
Binding of antibodies from CSF to live cells. (**A**) Binding of antibodies from CSF to astrocytes or neurons derived from human iPSC. Each column scatter plot shows results for CSF samples from donors with a diagnosis of CIS, clinically definite MS, or controls. Upper row of plots shows binding to neurons, and lower plots binding to astrocytes. Plots on left show IgM, plots in center IgG, and plots on right show IgA. Each dot represents one sample. Vertical axis shows geometric mean fluorescence intensity. Results from CSF samples from donors with CIS are shown in black, from MS in red, and from controls in blue. (**B**) Exemplary flow cytometry dot plot comparing IgM binding from CSF against the indicated cell lines. The vertical axis shows the fluorescence intensity of the secondary antibody used to detect the bound antibody, and the horizontal axis shows the intensity of the fluorescent vital dye Cell Trace Violet used to label the various cell lines before mixing, to enable their subsequent separation. Red dot plot shows IgM binding from CSF donated by a patient with CIS. Blue dot plot shows results with secondary antibody only. (**C**) Relationship between PNET-binding IgM signal (vertical axis, GMFI ratio) and CSF/serum IgM quotient (horizontal axis) (Pearson r = 0.096, p = 0.176). (**D**) Binding of antibodies from CSF to various cell lines. Vertical axis shows GMFI ratio (antibody+secondary-labelled cells divided by cells labeled with secondary antibody only). The 18 columns represent binding of IgM (upper row) or IgG (lower row) from CSF samples from donors in Basel with CIS (black), MS (red), and controls (blue), to each of six cell lines whose names are shown inside each plot. (one-way ANOVA * p = 0.049, **** p<0.0001). (**E**) Binding of antibodies from serum to a subset of the cell lines shown in (D). Vertical axis shows GMFI ratio (antibody+secondary-labelled cells divided by cells labeled with secondary antibody only). The 9 columns represent binding of IgM (on left) or IgG (on right) from serum samples from donors in Basel with CIS (black), MS (red), and controls (blue), to each of three cell lines whose names are shown inside each plot. (**F**) Binding of antibodies from CSF to the three cell lines shown in (E). Vertical axis shows GMFI ratio (antibody+secondary-labelled cells divided by cells labeled with secondary antibody only). The 9 columns represent binding of IgM (on left) or IgG (on right) from CSF samples from donors in Graz with CIS (black), MS (red), and controls (blue), to each of three cell lines whose names are shown inside each plot. (one-way ANOVA **** p<0.0001). (**G**) Binding of antibodies from serum to the three cell lines shown in (E). Vertical axis shows GMFI ratio (antibody+secondary-labelled cells divided by cells labeled with secondary antibody only). The 9 columns represent binding of IgM (on left) or IgG (on right) from serum samples from donors in Graz with CIS (black), MS (red), and controls (blue), to each of three cell lines whose names are shown inside each plot.

IgM binding varied both by disease status and by cell line (Fig. 1B-D). In particular, cells from the PNET and STTG1 lines were bound by IgM from CSF of patients with either CIS or MS, but not by IgM from control CSF (Fig. 1B-D). The PNET cell line was recognised by IgM from CSF from approximately 10% of MS patients and less than 1% of control samples, and this signal was not correlated with intrathecal IgM concentration –even among controls with relatively high levels of intrathecal IgM production, with one exception, no PNET-binding IgM was detected (Fig. 1C). CSF from six donors with MS contained cell-binding IgG, but this was discovered to result from therapeutically administered natalizumab, as recently described^14^, and these samples were therefore excluded from further analysis. Otherwise, CSF IgG was not different between patients with MS and controls (Fig. 1D).

For a subset of cell lines, including PNET, we examined binding of IgM and IgG from sera from the same donors. Both were similar across cell lines, and between MS and controls (Fig. 1E).

To confirm the MS-specific IgM binding pattern observed in CSF from the Basel cohort, we repeated the study using CSF samples from a cohort from an independent center (University Hospital of Graz, Austria). Samples (MS/CIS = 100, control = 100) were shipped coded and analysed by investigators unaware of diagnoses. Results replicated those from the Basel cohort (Fig. 1F, G). IgM binding to the PNET cell line was more commonly observed in patients with MS/CIS diagnosis, (CIS>control, p<0.0001, one-way, ANOVA) and after exclusion of 2 donors with therapeutic natalizumab detected, no disease-specific pattern was seen for IgG. Looking at clinical characteristics, those donors with anti-PNET IgM in their CSF were on average younger than negative donors, had more cells in CSF, and were more likely to have oligoclonal bands (Supplementary Table 3).

### Cloning and characterisation of PNET-specific CSF IgM antibodies

We prospectively screened CSF from a total of 212 patients for PNET binding. From one donor whose CSF contained PNET-binding IgM (Supp Fig. 2A), 1678 CD19+ B cells were sorted and cultured (Supplementary Fig. 2B). Five single B cell supernatants contained PNET-binding IgM (Fig. 2A, Supp. Fig. 2C). Transcriptomes of these B cells were surveyed by RNA sequencing, and immunoglobulin gene sequences were cloned (Supplementary Table 4), and one antibody (B3) showed the PNET-specific binding pattern of the original CSF (Fig. 2B, C). To gain some insight into the phenotype of the B cell that produced the B3 antibody, we compared its transcriptome with that of the other B cells that had undergone the same single cell transcriptomics protocol. B3 is characterised by an unusual combination of plasma cell markers such as PRMD1, IRF4, MZB1, GPR183 and JCHAIN, with genes related to antigen presentation such as HLA molecules, CD74 and CD53 (Fig. 2D, Supplementary Table 5). The 5 antibodies had various levels of somatic hypermutation (Supplementary Table 4). Following Beltran et al.^9^, we examined the ratio of silent to replacement mutations of the immunoglobulin gene. We found an increased rate of replacement mutations in the CDR of B3 (6.5) compared to the FR (1.4) indicating an antigen-driven mutation of this B cell clone.

**Figure 2.**
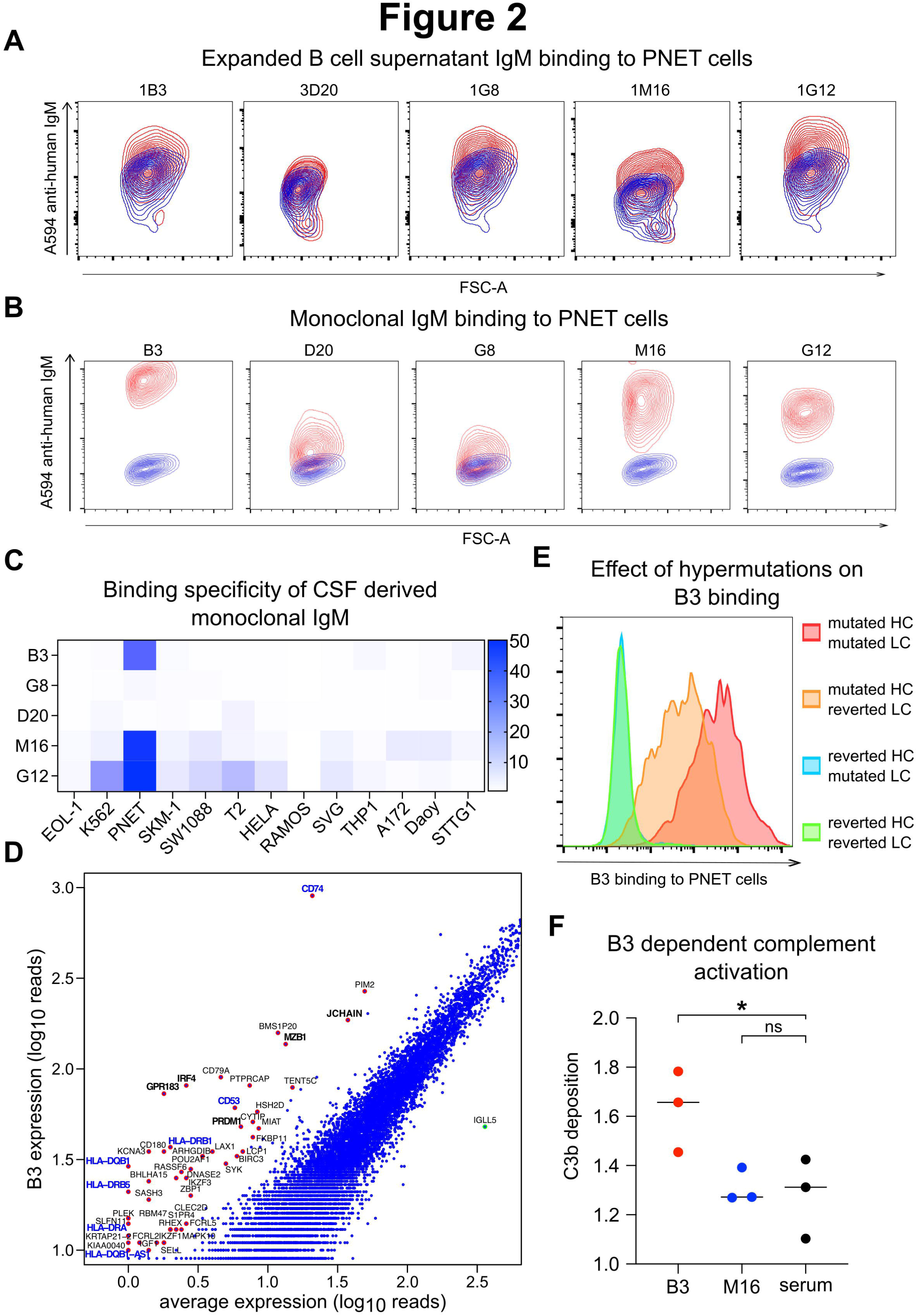
Properties of monoclonal IgM isolated from single CSF B cells. (**A**) Contour plots showing the PNET binding of IgM from the single cell culture supernatants from the wells highlighted in Supp. Fig. 2C. Names above each plot are the identities of the five wells, derived from their locations in the culture plates, and are used hereafter as identifiers for the monoclonal antibodies derived from these B cells. On each plot, the vertical axis shows IgM binding (fluorescence intensity of anti-human IgM), and the horizontal axis shows the forward scatter. Contours in blue are derived from cells labeled with anti-human IgM only, and contours in red from cells labeled with the single cell supernatants and then with anti-human IgM. (**B**) Contour plots like those shown in (A) representing binding of the five monoclonal antibodies cloned from the B cells in each of the five wells. (**C**) Heatmap showing binding of each of the five monoclonal antibodies (rows) to each of the 13 cell lines (columns). Blue intensity shown on bar at right corresponds to the GMFI ratio. (**D**) Transcriptional characteristics of B cell B3. cDNA from each of five singly cultured B cells was subjected to Illumina sequencing and for each gene observed, the log10 number of reads found in B3 was plotted against the log10 mean number of reads found in the other B cells (blue dots). Genes whose expression is greater than five-fold higher in B3 are marked with red circles and for a subset, their gene symbols printed beside the plotted dots. This information for all the B3 overabundant genes is collated in Supp. Table 5. Genes regarded as plasma cell markers have their symbols in black, genes associated with antigen presentation in dark blue. (**E**) Effect of mutational reversion on PNET binding. Binding of the recombinant antibody B3 with the observed mutations as originally cloned, in comparison with partially or fully germline-reverted versions of the same antibody. The red histogram shows binding of the original hypermutated antibody. The orange histogram shows binding of an artificial antibody with the native mutated heavy chain combined with a germline-reverted light chain. Blue histogram shows binding of antibody with reverted heavy and original mutated light. Green shows fully germline reverted version. (**F**) Antibody-dependent complement activation mediated by B3. PNET cells were incubated with monoclonal IgM B3, or with M16, in the presence of human serum from a healthy donor as a source of complement. Serum only without additional antibody was used as a control. Antibody-dependent complement activation was detected by bound C3 immunolabeling flow cytometry. Vertical axis shows GMFI ratio. *p = 0.0385, one-way ANOVA.

To further examine this inference, we tested the binding of a de-mutated version of the antibody, in which one or both of the chains were generated in their inferred germline sequences. Reversion of only the light chain partially reduced PNET binding, while reversion of the heavy chain completely eliminated it (Fig. 2E).

We also exploited complement activation as a measure of antigen-specificity. We observed that B3 was able to induce C3b deposition on the PNET cell surface (Fig. 2F) but the PNET-non-specific, and less mutated M16 was not.

### The molecular target of B3

B3 binding GMFI was reduced by roughly 10 fold following protease treatment (Fig. 3A), suggesting specific recognition of a cell surface protein. On the other hand, deglycosylation with various endoglycosylases did not affect the binding (Fig. 3A, Supp. Fig. 3). To directly investigate the antigenic target of B3, we used immunoprecipitation followed by mass spectrometry. This method has mostly been used with IgG, but artificially class-switching B3 to IgG compromised binding to PNET cells (Supp. Fig. 4A). We therefore adapted the technique for IgM using magnetic beads coated with anti-human IgM. We confirmed immunoprecipitation of haemagglutinin from transfected cells using a monoclonal anti-HA IgM (Supp. Fig. 4B)., and then applied the method to immunoprecipitate the target of B3 from PNET cells. Three independent experiments with a total of seven replicates identified several differentially retrieved proteins, and when these results were restricted to only those with the gene ontology (GO) term “plasma membrane”, the most obvious candidate was SCARA5 (Fig. 3B, Supp. Fig. 4C). In addition to the immunoprecipitation approach, we also reasoned that the specificity of the B3 antibody for the PNET cell line ought to be reflected in PNET-specific expression of the antigen, and we accordingly examined the whole transcriptome of each of the 13 cell lines, including PNET cell and the 12 non-bound cell lines (Fig. 3C). Several dozen genes were almost exclusively expressed in PNET cells and narrowing the search to plasma membrane proteins significantly more highly expressed in PNET than other cell lines yielded 30 candidates, including SCARA5 (Fig. 3C).

**Figure 3.**
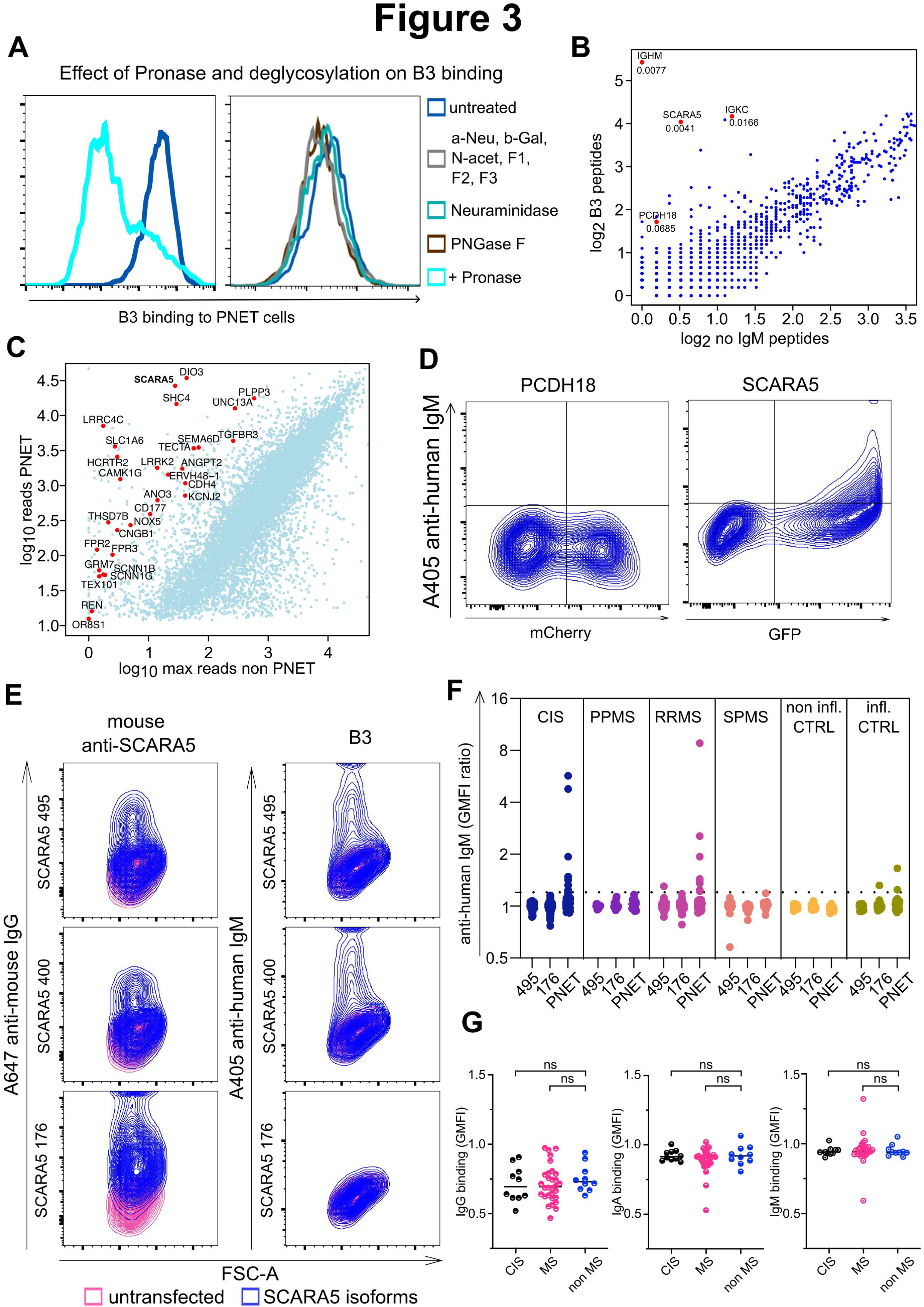
Target Antigen of IgM B3. (**A**) Effects of protease treatment or deglycosylation on B3 binding. Histogram on left shows binding to untreated PNET cells in dark blue, and binding to cells treated with the non-specific protease mixture Pronase to cleave exposed cell surface proteins in light blue. Histogram on right shows the effect of deglycosylation treatment on B3 binding. Binding to untreated PNET cells is shown in dark blue, binding to cells treated with various deglycosylating enzymes to specifically cleave terminal sugar moieties from cell surface carbohydrates is shown with other coloured histograms as specified on right of plot. Specificity controls for deglycosylation treatments are shown in Supp. Fig. 3. (**B**) Immunoprecipitation and mass spectrometry to identify the unknown protein target of the PNET-binding IgM B3. Live PNET cells were first incubated with B3, then washed and lysed; the lysate was next incubated with anti-IgM coated Dynabeads, washed and magnetically retrieved. Proteins were eluted from the beads, digested with trypsin, and peptides detected by mass spectrometry. Data are pooled from three independent experiments. Vertical axis shows log2 number of peptides precipitated by B3. Horizontal axis shows same parameter from control preparation in which the B3 was omitted. Each dot is one protein. Red dots are proteins whose retrieval was significantly more likely in the B3 samples (2-tailed unpaired t-test, p < 0.1), filtered for only those genes whose gene ontology (GO) terms include “plasma membrane”. These proteins are labeled with their gene symbol, and the p-value is shown below each symbol. (**C**) Transcriptional characteristics of the PNET cell line bound by B3. cDNA from each of 13 cell lines was subjected to Illumina sequencing and for each gene observed, the log10 number of reads found in the PNET cell lines was plotted against the log10 maximum number of reads found in the other B cell lines (light blue dots). Genes whose expression is significantly higher in PNET (2-tailed, unpaired t-test, p < 10-8), and whose GO terms include “plasma membrane” are marked with red circles and their gene symbols printed adjacent to the plotted. (**D**) Binding of B3 to cells transfected with either PCDH18 or SCARA5 measured by flow cytometry. B3-non-binding HEK cells were transfected with plasmids mediating expression of either PCDH18 (left) or SCARA5 (right), and co-transfected with fluorescent protein transfection markers (mCherry for PCDH18 and GFP for SCARA5). Vertical axis of each contour plot shows B3 binding, and horizontal axis shows transfection marker. (**E**) Impact of isoform. Analogous experiments to D were conducted using HEK cells transfected with each of the 176-, 400-, and 495 amino acid isoforms of SCARA5. Vertical axis shows binding of a commercial mouse anti-human SCARA5 antibody (left plots), or B3 (right plots). Horizontal axis shows forward scatter (no transfection marker used in this experiment). Blue contours are results with the commercial or B3 antibody, red contours are results from cells incubated with the appropriate secondary antibody only. (**F**) Prevalence of IgM antibodies against PNET cells and against SCARA5 in a cohort of donors with various diagnoses. Vertical axis shows IgM binding. Six sections left to right show results from CSF from donors with different diagnoses (left to right: CIS, PPMS, RRMS, SPMS, healthy controls, inflammatory non-MS, as specified within each plot). Within each of the six plots, the left-most column shows binding to HEK cells transfected with the 495-amino acid isoform, the center column binding to the 176-amino acid isoform, and the right column binding to PNET cells. (**G)** Binding of antibodies from serum to SCARA5-expressing cells. Vertical axis shows GMFI ratio (antibody + secondary-labelled cells divided by cells labeled with secondary antibody only obtained from SCARA5-HEK cells, again divided by the same value obtained from untransfected HEK cells). Each separate plot represents binding IgG (on the left), IgA (in the middle) or IgM (on the right), from serum samples from donors with CIS (black), MS (red) and controls (blue). Comparison of antibody binding between different diagnosis was performed by one-way ANOVA.

From these results, two genes appeared as plausible candidates, the iron transport scavenger protein SCARA5, and the protocadherin PCDH18. Flow cytometry with transiently transfected cells showed no specific binding of B3 to PCDH18-transfected cells, but clear binding to the SCARA5 transfected cells (Fig. 3D). PNET cells express three isoforms of SCARA5, with 495, 400, and 176 amino acids, and we repeated the flow cytometry screen with each. The two longer isoforms were clearly bound, while isoform 176 had binding no higher than untransfected cells (Fig. 3E). We confirmed the antigenicity of the transfected SCARA5 with commercially available antibodies and tested whether the epitope bound by B3 overlapped with these antibodies (Supplementary Table 6). The commercial monoclonal and polyclonal antibodies, and B3, all had individual, non-overlapping epitopes.

### Antigen validation and cohort screening

To test whether the newly identified antigen explains the PNET IgM reactivity we observed in the CSF of MS patients, we screened a cohort of 307 CSF samples (87 from RRMS, 116 from CIS, 23 from SPMS, 14 from PPMS, 27 from inflammatory controls, 40 from non-inflammatory controls) for the presence of antibodies binding to PNET cells, or to HEK cells transfected with 495– or 176-amino acid isoforms of SCARA5. We confirmed the binding of CSF IgM to PNET cells in 11 % of CIS, 10 % of RRMS patients and 3% of control patients. In one RRMS case, IgM binding to PNET cells also bound to SCARA5 isoform 495. None of the PPMS, SPMS or non-inflammatory controls showed reactivity against PNET cells or cells expressing SCARA5 isoform 495 (Fig. 3F). We also tested the serum reactivity against SCARA5-transfected HEK cells for a subset of donors, and found only one donor (distinct from the donor with anti-SCARA5 IgM in CSF) with detectable anti-SCARA5 IgM in serum (Fig. 3G).

For a subset of donors, we compared the anti-PNET IgM signal with the abundance of paramagnetic rim lesions, as assessed by MRI, but there was no significant difference in this parameter between positive and negative donors (Supplementary Table 3).

### Anti-SCARA5 antibodies influence leukocyte trafficking into the CNS

To investigate possible mechanisms by which antibodies against SCARA5 might influence the autoreactive immune response, we examined the expression of SCARA5 in autoptic brain lesions from MS donors by immunolabeling with a rabbit polyclonal anti-SCARA5 antibody. Most intense labeling was observed on foamy macrophages adjacent to active lesions, but weaker labeling was also found associated with neurons, and astrocytes around chronic MS lesions (Fig. 4A, Supp. Fig. 5). Occasionally, SCARA5 labeling was overlapping with IgM deposition in MS lesions (Fig. 4B). To corroborate these results, we interrogated publically available databases of single cell RNA expression data bioinformatically. This approach showed expression of SCARA5 RNA in gabaergic interneurons, mesenchymal cells of the choroid plexus, and endothelial cells (Fig. 4C). Immunolabeling of mouse brains showed strongest expression in the periventricular and sub-dural blood vessels, and in the choroid plexus (Fig. 5A). In view of the importance of blood vessels and the choroid plexus in immune cell trafficking into the CNS^14^ we hypothesized that anti-SCARA5 antibodies might exert their pathogenic effect by perturbing immune cell migration. We confirmed that anti-SCARA5 antibodies injected directly into the lateral ventricle bind to periventricular blood vessels in vivo (Fig. 5B), and then examined the effect of this manipulation on immune cell trafficking (Fig. 5C). We transferred activated 2D2 T cells, specific for the myelin protein myelin oligodendrocyte glycoprotein (MOG) into mice, and as the first animal manifested motor signs, we injected half of the animals with polyclonal sheep anti-SCARA5 antibody (the patient-derived antibody B3 is human-specific), or control antibody (sheep anti-influenza neuraminidase) into the lateral ventricle. We tracked the weights and motor behavior of the animals (Fig. 5D, E), and one week later we sacrificed the animals and quantified T cells in the brain by immunofluorescence microscopy. T cell infiltration in the perivascular areas adjacent to the injection site was significantly augmented by anti-SCARA5 antibodies (Fig. 5F, G).

**Figure 4.**
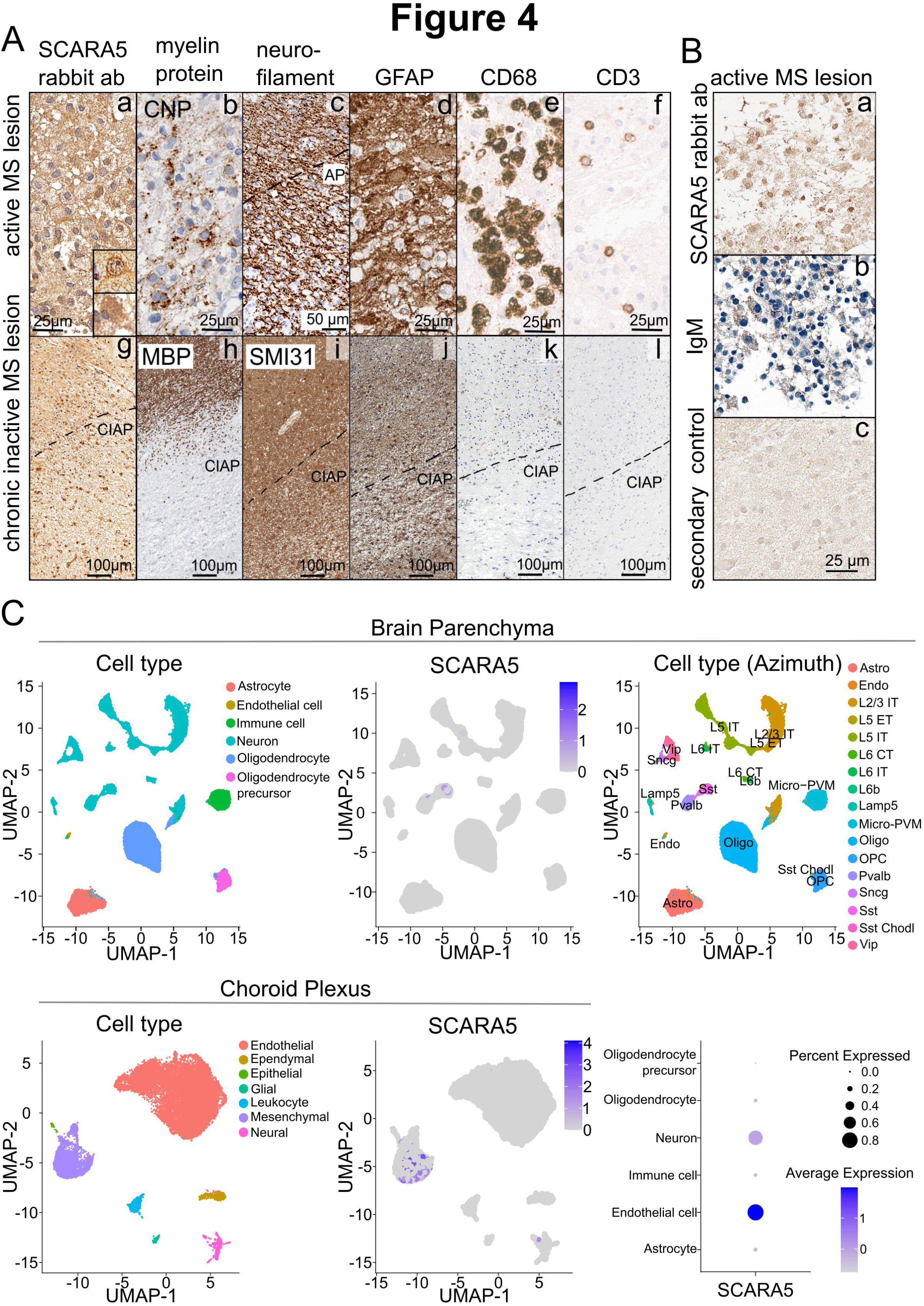
Immunohistochemical localisation of SCARA5 in human brain. (**A**) In a-f, the immunohistochemical staining of an early active MS lesion is shown. In **a**, in particular, the numerous macrophages are positively labeled with SCARA5-antibody. In addition, some astrocytes are positively labeled in this staining (lower inset). In the vicinity of the lesion in the cortex, some neurons are labeled, one of which is shown in the upper inset in a. In the staining against the myelin protein CNP-ase (CNP) in **b**, some macrophages with CNP-positive myelin degradation products are shown within the lesion. Neurofilament heavy chain staining in **c** shows the rarefied but preserved axonal scaffold within the lesion (lesion boundary corresponds to dashed line, AP = active plaque). The GFAP staining (**d**) highlights the marked reactive astrocytosis. In the staining against CD68 (**e**), the numerous (foamy) macrophages are positively labeled. In (**f**), the sparse T-lymphocytic infiltrate is emphasized in staining against CD3. (**g-l**) In g-l, the immunohistochemical staining of a chronic inactive MS lesion is shown. In the first picture (**g**), activated microglial cells and reactively altered astrocytes are positively labeled in the inactive lesion stained with SCARA5 (lesion boundary corresponds to dashed line, CIAP = chronically inactive plaque). Myelin sheath immunostaining against the myelin protein MBP shows the sharp lesion boundary (**h**). In the staining against SMI31 (phosphorylated neurofilament heavy chain), the rarefied but preserved axonal scaffold is shown within the lesion (**i**). In J, the marked reactive astrocytosis is shown in the GFAP staining. Reactive microglial cells are positively labeled in k (CD68). No T lymphocytes are present in this lesion (CD3 in **l**). **B (a-c)** An additional early active MS lesion is labeled with SCARA5 antibody (a) and IgM deposition is concomitantly detected (b). **C** Visualization of cell type distribution and relative SCARA5 expression in single-nucleus transcriptomes from 30 frontal cortex and choroid plexus samples^29^ using uniform manifold approximation and projection (UMAP) in either data set and dot plot illustrating expression frequency and intensity. Cell type annotations are based on the original publication for general cell types and public reference-based mapping for neuronal subclasses in human cortex tissue (Azimuth), respectively.

**Figure 5.**
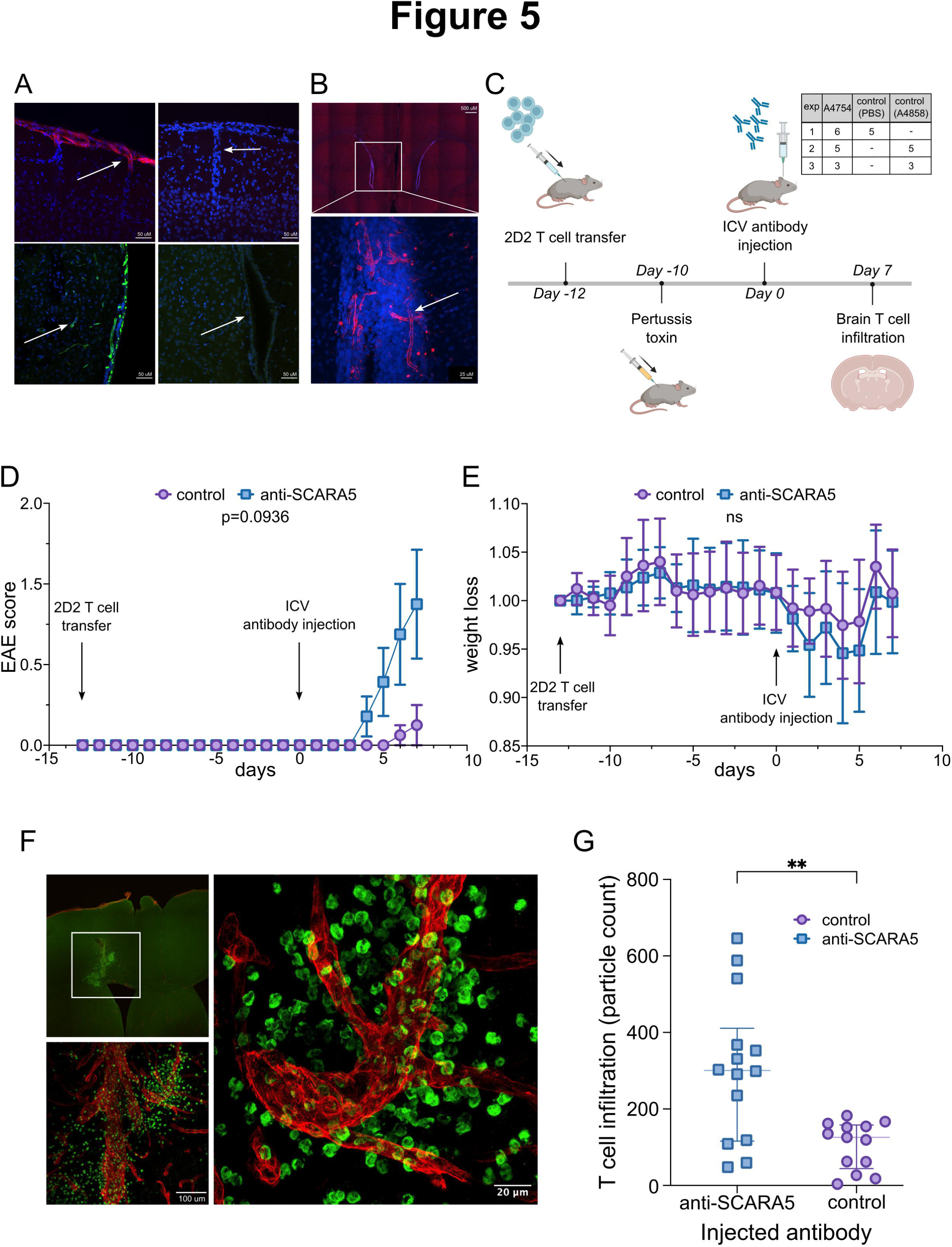
Impact of anti-SCARA5 antibodies on immune cell trafficking into brain. (**A**) Immunofluorescent labeling of SCARA5 expression in mouse brain. Vibratome sections from perfusion-fixed brains of unmanipulated mice were immunolabeled with sheep anti-SCARA5 antibodies and visualised by confocal microscopy. Two upper images show surface (sub-dural) and penetrating blood vessels of the superficial cortex. The upper left shows nuclei labeled with DAPI in blue, and immunolabeling for SCARA5 in red. The upper right shows a similar location from a control section, from which the primary antibody was omitted, imaged with the same parameters. In the lower two images, labeling of periventricular blood vessels is shown. The left image is labeled with sheep anti-SCARA5 (green) and the right is a no-primary control. (**B**) Anti-SCARA5 binding to live blood vessels. A polyclonal sheep anti-SCARA antibody was injected into the lateral ventricle of a mouse, and the next day the mouse was sacrificed by perfusion fixation, and the brain labeled with DAPI (blue) and anti-sheep secondary antibody (red). The upper panel is a composite image made by stitching together images captured with a 10x objective, to show both the ventricles – the ventricle highlighted with a box on the left received the injection. The panel below is a maximum intensity projection of a confocal stack of the area shown in the box, captured with the 40x objective showing the blood vessels in the wall of the ventricle labeled by the in vivo antibody injection. (**C-G**) Impact of intraventricular anti-SCARA5 antibodies on encephalitic T cell infiltration. All figures show results pooled from 3 experiments. (**C**) Schematic diagram of experimental design (created with BioRender.com) and table of animal numbers per experiment. In vitro activated, MOG-specific 2D2 T cells were transferred to recipient animals by intraperitoneal injection, followed by pertussis toxin, and animals were tracked until the first animals showed motor signs, at which point, the remaining asymptomatic animals were randomised into two groups to receive anti-SCARA5 antibody or control. (**D**) Motor impairment of animals over time. Motor function was subjectively assessed according to a system used for scoring experimental autoimmune encephalomyelitis (EAE), in which 0 is unimpaired, and 4 is severe impairment. Points show the means within the conditions and the bars the standard error of the mean. Purple circles represent control animals, and blue squares the animals injected with anti-SCARA5. p=0.0936 calculated by two-tailed unpaired t-test between the areas under the curves for each condition. (**E**) Weights of animals over time of experiment. Weights are expressed as the weight at each time point divided by the weight at the start of the experiment for each animal. Points show the means within the conditions and the bars the standard error of the mean. Purple circles represent control animals, and blue squares the animals injected with anti-SCARA5. (**F**) T cell infiltration. Vibratome section of brain from animals sacrificed at one week after intraventricular antibody injection in the experiment described above were immunolabeled for laminin (blood vessel walls, red) and CD3-epsilon (T cells, green). (**G**) Quantification of infiltrating T cells. Images like the ones shown in F were captured for 5-6 sections per mouse by an automated fluorescent microscope, and the numbers of T cells per section extracted by an automated script in ImageJ. Points show the median number of T cells per section for each mouse and the whiskers show the interquartile range. ** p = 0.0012, two-tailed unpaired t-test.

## Discussion

Antibodies are produced intrathecally in several CNS disorders, and OCB are found in the CSF of more than 90% of patients with MS^3,8^. The most obvious outstanding question about CSF antibodies in MS concerns their target antigen(s).

In this study, we aimed to identify the target of intrathecally produced immunoglobulins by screening them against multiple neuronal cell lines and human iPSC-derived neurons and astrocytes. At the population level, we identified an IgM signal in CSF that discriminated MS patients from controls. From a single B cell of a patient whose CSF showed this signal, we isolated a monoclonal antibody and identified its molecular target as the iron transporting scavenger protein SCARA5.

Most investigations of intrathecally produced antibodies in MS have focussed on IgG, however, intrathecal IgM synthesis is associated with a more active inflammatory disease phenotype^6,15^, a higher likelihood of conversion from CIS to clinically definite MS^7,16–18^, and spinal manifestation, suggesting a distinct clinical phenotype and pathophysiology^8,19^. Intrathecal IgM is also associated with increased complement activation in the CSF, especially of the early classical and alternative pathways (Oechtering et al., under revision). This observation implies that the IgM must find antigen in the CNS, because complement activation by IgM in the classical pathway requires a multimeric cognate interaction between multiple Fab domains and their targets ^20^.

Additional evidence for an antigen-driven intrathecal origin of the anti-PNET IgM includes the fact that B3, like the IgM antibodies described by Beltrán et al^9^ shows CDR-enriched somatic hypermutation, and that antigen-binding is dependent on this mutation.

The transcriptional profile of the B cell giving rise to B3 was partially typical of a plasma cell. The high expression of HLA genes seen in this B cell is also consistent with a plasma cell phenotype and has been associated with freshly generated, antigen-specific plasma cells thought to be pathogenic in systemic lupus erythematodes^21^. High expression of GPR183 by B3 was unexpected. GPR183 is thought to be involved in directing B cell migration within lymph nodes^22^, and has more recently been shown to contribute to tissue homing and antibody secretion by plasma cells in the gut^23^.

SCARA5 functions as a transferrin-independent ferritin receptor for both iron delivery and ferritin removal ^24,25^. Dysregulation of iron metabolism and the resultant cytotoxicity is increasingly implicated in MS^26^. Elevated transcripts of SCARA5 as well as of other iron import related molecules were described in MS lesions, especially in the periplaque white matter, suggesting a cellular defense response against iron toxicity ^27^. Soluble SCARA5 was also significantly reduced in the CSF of patients with MS compared to controls^28^. Considering the expression of SCARA5 in brain endothelial cells and choroid plexus we hypothesized that anti-SCARA5 antibodies could exert their pathogenic effect by influencing immune cell trafficking into the brain. Intrathecal application of a SCARA5 antibody had indeed an influence on T cell infiltration in an EAE model using MOG specific T cells.

Since the proportion of patients with anti-PNET IgM in CSF is much larger than the population with anti-SCARA5 reactivity, there must be a large population of patients with antibodies binding some other epitope on the PNET cells. One possibility is that SCARA5 is one protein in a complex that is the target of the MS-specific antibody response. However, SCARA5 is not an intensely studied protein, and its binding partners have yet to be characterized, so additional studies will be required to investigate this possibility.

### Data availability

All data produced in this study will be made publicly available by the time of publication with the exception of human sequence data with the potential to infringe donors’ rights to withdraw consent for their data to be made public. In particular, the complete coding sequences of heavy and light chains of the antibodies described can be found at ncbi accession number (accession number OQ744174.1 to OQ744183.1).

**Abbreviations:** CIS = clinically isolated syndrome; CTRL = control; GO = gene ontology; IgM = immunoglobulin M; iPSC = induced pluripotent stem cells; MS = multiple sclerosis; OCB = oligoclonal bands; PNET = primitive neuroectodermal tumor

## Supporting information

Supplementary Figure Ledends

## Acknowledgements

We thank the patients and staff of the University Hospital Basel and the University Hospital of Graz for their participation in the study. We thank Barbara Treutlein and Ryoko Okamoto for the provision of cells for testing antibodies. We thank Rachel Lindemann, for the support in conducting some of the animal experiments. Sample collection was made possible by the Swiss Multiple Sclerosis Cohort organisation, supported in part by the Swiss Multiple Sclerosis Society. We are also grateful for technical support from the flow cytometry, microscopy, and bioinformatics core facilities of the Department of Biomedicine, and computational resources provided by the scientific computing center of the University of Basel (http://scicore.unibas.ch/).

## Funding

The authors disclosed receipt of the following financial support for the research, authorship, and/or publication of this article: Funding was provided by the Swiss Multiple Sclerosis Society (Nicholas Sanderson) and by the Swiss National Science Foundation grant number 189043 (Tobias Derfuss). Ilaria Callegari received funding from the European Academy of Neurology. Martin Diebold received funding from the Swiss National Science Foundation [project-number 199310] and the German Research Foundation [IMM-PACT-programme, 413517907].

### Supplementary material

Supplementary material is available at *Brain* online.

