## Supplementary Methods for "Cell-binding IgM in CSF is distinctive of multiple sclerosis and targets the iron transporter SCARA5"

### CSF single B cell expansion and immunoglobulin sequencing

B cells from PNET-binding IgM positive CSF samples were sorted and expanded using the protocol described by Huang et al. [1] and subsequent modifications. Briefly, as soon as a CSF sample arrived, CSF cells were resuspended R40, transferred to a sterile flow cytometry tube, spun down at 300 x g, at 4°C for 3 minutes and resuspended in 100µlof PBS plus 1µlof the following antibodies: FITC-conjugated anti-CD3 (BioLegend 317306), anti-CD4 (BioLegend 357406), anti-CD14 (BioLegend 301804), anti-CD16 (BioLegend 360716), anti-CD56 (BioLegend 318304); PE-conjugated anti-CD19 (BioLegend 363004). Cells were incubated on ice for 20 minutes, then washed once, resuspended in 100µlof sterile PBS and sorted using a FACS Aria III (BD Biosciences). The gate was set on the CD19-positive, CD3/CD4/CD14/CD56-negative population. Sorted CD19+ cells were cultured in single 384-well plate wells at an average density of 0.9 cells/well, in R40, 0.05 ng/µl IL-21 (Gibco, PHC0215), together with 50,000 irradiated (75 Gy) TE CD40L cells/well (Edgar Meinl, Ludwig-Maximilians-Universität, Munich, Germany). After 9 days of culture, 15 µl of supernatant was withdrawn from each well and screened for PNET-binding antibodies as described above. B cells from positive wells were lysed in 20 µl of 15 mM Tris–HCl, pH 8.0 containing 0.5 U/µl of recombinant murine RNAse inhibitor (NEB, M0314L). RNA was extracted from this lysate by Zymo Quick-RNA microprep kit (R1050 & R1051), and reverse transcribed with the SMART-Seq v4 Ultra Low Input RNA Kit for Sequencing (Takara, 634888). One hundred ng of the thus generated cDNA was used to prepare sequencing libraries with the DNA prep kit (Illumina). The libraries were sequenced paired-end 150 bp on a NextSeq 500 sequencer (Illumina) yielding an average of one million reads per lysate. Reads mapping to immunoglobulin gene sequences were extracted from the resulting single-cell transcriptomes with a custom script in R (Dataset EV3 and 4), using packages Shortread [2] and Biostrings, reassembled with EZassembler [3], and assigned to V(D)J genes using IgBLAST (https://www.ncbi.nlm.nih.gov/igblast/).

### Recombinant antibody production

Based on the deduced heavy and light chain sequences, DNA constructs encoding the inferred amino acid sequence from leader to several bases into the constant region were synthesized by

IDT with restriction sites in the termini to enable in-frame cloning into pVITRO (Invivogen) and the resulting plasmids transfected into Freeestyle HEK cells (Invitrogen, R79007). On the day of transfection, each 125 ml flask containing 14 ml of cells at 1 x 10^6 viable cells/mL was transfected with pVITRO hygro dual expression plasmid expressing the heavy and the light chain of each of the recombinant antibodies, using 293fectin Transfection Reagent (Gibco, 12347019) according to the manufacturer's instructions. 24 h post-transfection, hygromycin B (50 μg/ml) was added to transfected cells, which were then maintained in culture under selection for 2 weeks at densities between 3 x 10^5 and 5 x 10^5 viable cells/ml. Cultured supernatants were harvested after 16 days, centrifuged at 2000 g for 20 minutes, passed over 0.45 mm filters (Sartorius). Cell supernatants were buffer-exchanged into PBS using Amicon Ultra 15 50-K columns (Sigma, UFC905024) following the manufacturer's instructions, and the resulting antibody containing supernatants were stored at 4°C or at -20°C until use. Recombinant antibodies were quantified by ELISA. For the preparation of antibodies with germline-reverted heavy and/or light chains, germline sequences corresponding to the cloned V heavy and V light genes were obtained from IMGT (https://www.imgt.org/), synthesized and cloned into pVITRO exactly as for the mutated sequences. Antibodies were then produced with either both germline chains, both mutated, or one germline and one mutated, by switching the coding regions in pVITRO.

### Differentiation of human induced pluripotent stem cell (hiPSC)-derived neurons and astrocytes

Human iPSCs were generated from peripheral blood mononuclear cells (PBMCs) of one healthy donor (Male donor age 50) as previously described[4]. The differentiation into neurons and astrocytes has been detailed other publications[5,6]. Briefly, hiPSCs were first differentiated into NPCs as per this detailed protocol. For neuronal differentiation, NPCs were seeded on poly-L-ornithine/Laminin-coated plates at 50,000 cells/cm$^2$ in Neural medium (DMEM/F12 + 1x N2 supplement, 1x B27 supplement, 2x Culture One supplement, 2 μg/ml Laminin, 20 ng/ml BDNF). The medium was renewed every 2-3 days for 14 days before using the neurons for the assay. For astrocyte differentiation, NPCs were plated onto matrigel-coated dishes in DMEM/F-12 supplemented with 1x N2 supplement, 1x B27 supplement, 20ng/ml LIF and 10ng/ml EGF for two weeks, then in Astrocyte medium (DMEM/F-12 with 1x B27 supplement) supplemented with 20 ng/ml CNTF for another 4 weeks. When mature, astrocytes were used afterwards for downstream assays.

### Flow cytometry

For flow cytometry screening of PNET-binding antibodies in CSF, 100000 PNET cells and the same number of SW1088 cells per well were incubated with a 1 to 4 dilution of the CSF in PBS. Incubation was done in 96-well-plates, for 30 minutes, on ice. Cells were then washed twice with cold PBS and labeled with 100 µl of PBS containing Dylight-405-conjugated anti-human IgM (JIR 109-475-129), FITC-conjugated anti-human IgG (JIR 109-096-098) and Alexa Fluor 647-conjugated anti-human IgA (JIR 109-605-011) for 30 minutes on ice, washed twice with cold PBS and measured on a Beckman Coulter CytoFLEX flow cytometer equipped with a 96-well plate reader.

A similar technique was used for testing the supernatants from expanded B cells; in this case, 10 µl PNET cells were incubated on ice with 15 µl of supernatant from each expanded B cell, in a 384-well plate (Thermo Scientific). After 30 minutes' incubation on ice, cells were transferred to a 96-well plate, washed twice in PBS and labeled as described above.

For testing effect of surface protein removal on monoclonal antibody binding, cells were trypsinised, washed twice with PBS and resuspended in HBSS 5 mM CaCl plus Pronase (SigmaAldrich) 2 mg/ml, then incubated for 1h at 37°C. Cells were then labeled with each monoclonal antibody or CSF as described above. Deglycosylation used PNGase, O-Glycosidase, α-2(3,6,8,9)-Neuraminidase, Endoglycosidase F1, F2, F3 alone or in combination. Cells were incubated at 37 °C for 3 hours with each enzyme, washed and incubated with each patient-derived or control antibody or lectin as described in the figure legends.

To assess complement activation by patient-derived antibodies, we incubated PNET cells with antibodies at 37°C for 3 hours with 20% human serum as a complement source, then washed them and detected the bound C3b component with a mouse monoclonal (Biolegend cat #846402) and a rhodamine RedX-conjugated anti-mouse secondary (Jackson115-295-166) by flow cytometry.

### ELISA

384-well plates were coated with goat anti-human IgG (Southern Biotech, 2014-01), anti-human IgM (Southern Biotech, 2023-01), or anti-human IgA (Southern Biotech, 2053-01) antibodies overnight at 4°C, washed once with PBS and blocked with PBS- 1% BSA at room temperature for 90 minutes. Plates were then washed three times with PBS 0.05% Tween, incubated with 15 µl of serially diluted samples for 2 h at room temperature, washed three times with PBS 0.05% Tween and incubated with anti-IgG-HRP (Southern Biotech 2014-05), anti-IgM-HRP (Southern Biotech 2023-05), anti-IgA-HRP (Southern Biotech 2053-05) or goat anti-

Ig-HRP (Southern Biotech, 2010-05) in PBS-0.1% BSA for 1h at room temperature. Plates were then washed three times with 80 µl/well of PBS-0.05% Tween and developed with TMB ELISA substrate (SureBlue Reserve TMB Microwell Peroxidase Substrate, REF 53-00-00) until a blue color was visible; the reaction was stopped with sulphuric acid and plates were read at 450 nm immediately after stopping.

### Immunoprecipitation and mass spectrometry

Cultured PNET cells from three T150 flasks were washed twice with ice cold PBS and then incubated with recombinant B3 for 30 minutes at 37°C, washed twice in PBS and trypsinised. Cells were then retrieved in R10, centrifuged at 300 g for 3 minutes at 4°C, washed twice in PBS, and the pellet lysed in 500µlof Pierce IP lysis buffer containing Halt Protease Inhibitor Cocktail (Thermo Fisher). The lysate was sonicated at low power for 10 1 second pulses and incubated on ice for 30 minutes, then centrifuged for 30 minutes at 10’000 g at 4°C. The supernatant was then incubated with Dynabeads (Thermo Fisher, 5 mio of beads every 40 mg of cell pellet) previously coated with unconjugated Donkey anti-human IgM (JIR 709-005-073) for 30 minutes at 4°C on a rotor. The suspension was then washed twice with Pierce IP lysis buffer. The product from immunoprecipitation was eluted by incubating the suspension with 106 mM Tris HCl, 5% LDS and boiling the samples for 10 min at 70°C. The elution product was run on an 4-12% Bis-Tris gels (Thermo Fisher) in MOPS running buffer, transferred to a nitrocellulose membrane and probed by immunoblot using Alexa Fluor 680 conjugated streptavidin to detect biotinylated membrane proteins and each our recombinant IgM. Samples were then cooled down to RT and 0.5 µL of 1M iodoacetamide was added to the samples. Cysteine residues were alkylated for 30 min at 25°C in the dark. Digestion and peptide purification was performed using S-trapTM technology (Protifi) according to the manufacturer’s instructions. In brief, samples were acidified by addition of 2.5 µL of 12% phosphoric acid (1:10) and then 165 µL of S-trap buffer (90% methanol, 100 mM TEAB pH 7.1) was added to the samples (6:1). Samples were briefly vortexed and loaded onto S-trapTM micro spin-columns (Protifi) and centrifuged for 1 min at 4000 g. Flow-through was discarded and spin-columns were then washed 3 times with 150 µL of S-trap buffer (each time samples were centrifuged for 1 min at 4000 g and flow-trhough was removed). S-trap columns were then moved to the clean tubes and 20 µL of digestion buffer (50 mM TEAB pH 8.0) and trypsin (at 1:25 enzyme to protein ratio) were added to the samples. Digestion was allowed to proceed for 1h at 47 °C. After, 40 µL of digestion buffer was added to the samples and the peptides were collected by centrifugation at 4000 g for 1 minute. To increase the recovery, S-trap columns were washed with 40 µL of 0.2% formic acid in water (400g, 1 min) and 35 µL of 0.2% formic acid in 50% acetonitrile. Eluted peptides were dried under vacuum and stored at -20 °C until further analysis.

### Transcriptomic analysis

RNA was extracted from pelleted cell lines with TriZOL accordig to the manufacturer's instructions and quality-checked on the TapeStation instrument (Agilent Technologies, Santa Clara, CA, USA) using the RNA ScreenTape (Agilent, Cat# 5067-5576 - Average RINe 9.8±0.3) and quantified by Fluorometry using the QuantiFluor RNA System (Cat# E3310, Promega, Madison, WI, USA).

Libraries were prepared from 70 ng total RNA per cell line, using the TruSeq Stranded mRNA Library Kit (Cat# 20020595, Illumina, San Diego, CA, USA) and the TruSeq RNA UD Indexes (Cat# 20022371, Illumina, San Diego, CA, USA) with 15 cycles of PCR and quality-checked on the Fragment Analyser (Advanced Analytical, Ames, IA, USA) using the Standard Sensitivity NGS Fragment Analysis Kit (Cat# DNF-473, Advanced Analytical). Average concentration was 83±13 nmol/L and average library size was 329±6 base pairs. Samples were pooled to equal molarity andquantified with the QuantiFluor ONE dsDNA System (Cat# E4871, Promega, Madison, WI, USA). Pooled libraries were sequenced Paired-End 38 bases (in addition: 8 bases for index 1 and 8 bases for index 2) using the NextSeq 500 High Output Kit 75-cycles (Illumina, Cat# FC-404-1005) loaded at 2.0pM and including 1% PhiX. Primary data analysis was performed with the Illumina RTA version 2.4.11 and Basecalling Version bcl2fastq v2.20.0.422. Two Nextseq runs were performed to compile enough reads (on average per sample: 507±15.5 millions pass-filter reads). Reads were aligned to the human genome and quantified using the QuASR package in R[7].

### SCARA5 and PCDH18 cloning

RNA was extracted from PNET cells using quickRNA (Zymo) following the manufacturer's instructions and reverse transcribed using SuperScriptTM III RT (200 units/µl), followed by removal of RNA by incubating the cDNA with E.coli RNase H at 37°C for 20 minutes. SCARA5 and PCDH18 sequences amplified by nested PCR and the resulting products cloned into the vPigLIC expression vector for transient transfection of HEK293 cells.

### Neuropathology and immunohistochemistry

The material from the autopsied MS patient was obtained from the Netherlands Brain Bank [NBB], where it was evaluated and the diagnosis confirmed by Andreas Junker. The MS biopsy material was obtained from the archive of the Institute of Neuropathology of the University Hospital Essen where it had been diagnosed. The study was approved by the ethics committee of the Ethics Commission of the University of Duisburg-Essen (reference: 16-6933-BO). All investigations were performed in compliance with relevant laws and institutional guidelines, and were carried out following the rules of the Declaration of Helsinki of 1964, revised in 2013. All investigations were performed on 1 μm sections. In addition to standard staining with hematoxylin-eosin (HE) (not shown), immunohistochemical staining was performed with antibodies against SCARA5 (MAB4900, 1:50), CNP (Sternberger, SMI91, 1:200), MBP (DAKO, REF A0623, 1:2000), Neurofilament (Sigma, N0142, 1:400), CD68(DAKO, PG M1, 1:50, and CD3 (DCS, clone SP7, 1:50) according to standard procedures. Pretreatments were carried out with citrate buffer. The endogenous peroxidase activity was first blocked by incubation of the sections in 3% $H_2O_2$ in PBS. This was followed by a blocking step with 10% fetal calf serum in PBS for ten minutes at room temperature, followed by incubation with the primary antibody for two hours at room temperature. The sections were then incubated with the secondary antibody (biotin labelled antibody). Finally, the staining was developed with 3,3'-diaminobenzidine (DAB). Cell nucleus counterstaining was performed with hematoxylin.

### Paramagnetic rim lesion count

Three-dimensional echo planar imaging (3D-EPI) sequence was available and used to detect PRL lesions (PRLs). Two trained raters independently examined the presence of PRLs. Ultimately, the PRL count was established through a consensus agreement. PRLs were defined as distinct FLAIR-hyperintense lesions encircled, either entirely or partially, by a rim of paramagnetic signal on susceptibility-based images. To ensure the chronic nature of the identified PRLs, we only considered for assessment lesions that did not exhibit enhancement in a concomitant post-contrast T1 image or, in the absence of this acquisition, lesions visible in a FLAIR image acquired at least 6 months prior.

### Single-nucleus transcriptomes from brain parenchyma and choroid plexus

Single-nucleus transcriptomes from 30 publicly available frontal cortex and choroid plexus samples from a well-described cohort[8] were employed and re-assessed in combination with standardized cell type annotation tools (based on original publication (choroid plexus data set) and HuBMAP's public reference-based mapping (Azimuth, cortex data set[9]). The cortex data set was further assessed by sub-setting endothelial cells and neurons with assessment of SCARA5 expression patterns within subsets. Testing and visualization of the sequencing data was performed using the Seurat package [10] in the R programming environment.

### Mouse model of anti-SCARA5 pathomechanism

MOG-specific 2D2 T cells were obtained from C57BL/6-Tg(Tcra2D2,Tcrb2D2)1Kuch/J ("2D2") mice. Recipient mice were B6.129P2(C)-*Cd19*$^{tm1(cre)Cgn}$/J x B6.Cg-*Gt(ROSA)26Sor*$^{tm14(CAG\text{-}tdTomato)Hze}$/J mice, congenic on the C57Bl/6 background, both bred at the University of Basel from founder animals obtained from Jackson. These recipient mice have B cells that express the red fluorescent protein tdTomato to facilitate B cell tracking by microscopy, but are otherwise similar to wild type C57Bl/6 mice. Splenocytes from 2D2 transgenic mice were cultured at 37°C in RPMI with 10% FCS, 50 µM 2-mercaptoethanol with penicillin and streptomycin (R-10) with 20 µg/ml MOG35-55 peptide (Anaspec), 5 ng/ml IL-2 and 5 ng/ml IL-7 (peprotech) for 24 hours, then in the same conditions for 4 days more but without MOG peptide, then finally stimulated with plate bound anti-CD3 anti-CD28 (Biolegend) for 24 hours before washing, counting, and transferring into recipient mice by intraperitoneal injection at 1.5 million cells per mouse. Two days later, each mouse received 66 ng of pertussis toxin (Millipore #516561) by intraperitoneal injection. Mice were weighed and scored daily, and when the first animal manifested any motor signs indicative of encephalomyelitis (weak tail), the remaining asymptomatic animals were randomized into two groups. One group received an intracerebral injection of 3 µl polyclonal sheep anti-SCARA5 antibody (BioTechne AF4754), and animals in the second group received 3 µl of a control antibody (BioTechne AF4858 sheep anti-influenza hemagglutinin) or PBS vehicle as specified in Figure 5B. Intracerebroventricular injections were performed under isofluorane anesthesia with a Hamilton syringe using a stereotactic frame (Stoelting). Scoring was done by investigators unaware of the experimental groups, by the following system: 0.5 noticeable tail weakness; 1 limp tail; 2 mild hind limb paresis; 2.5 strong hind limb paresis; 3 hind limb hemiplegia; 3.5 bilateral hind limb paralysis; 4 forelimb weakness. All procedures using live animals were reviewed and authorized by the Animal Research Commission of the Basel Cantonal Veterinary Office.
