## Supplementary figures and images for "Cell-binding IgM in CSF is distinctive of multiple sclerosis and targets the iron transporter SCARA5"

### Supplementary Figure 1

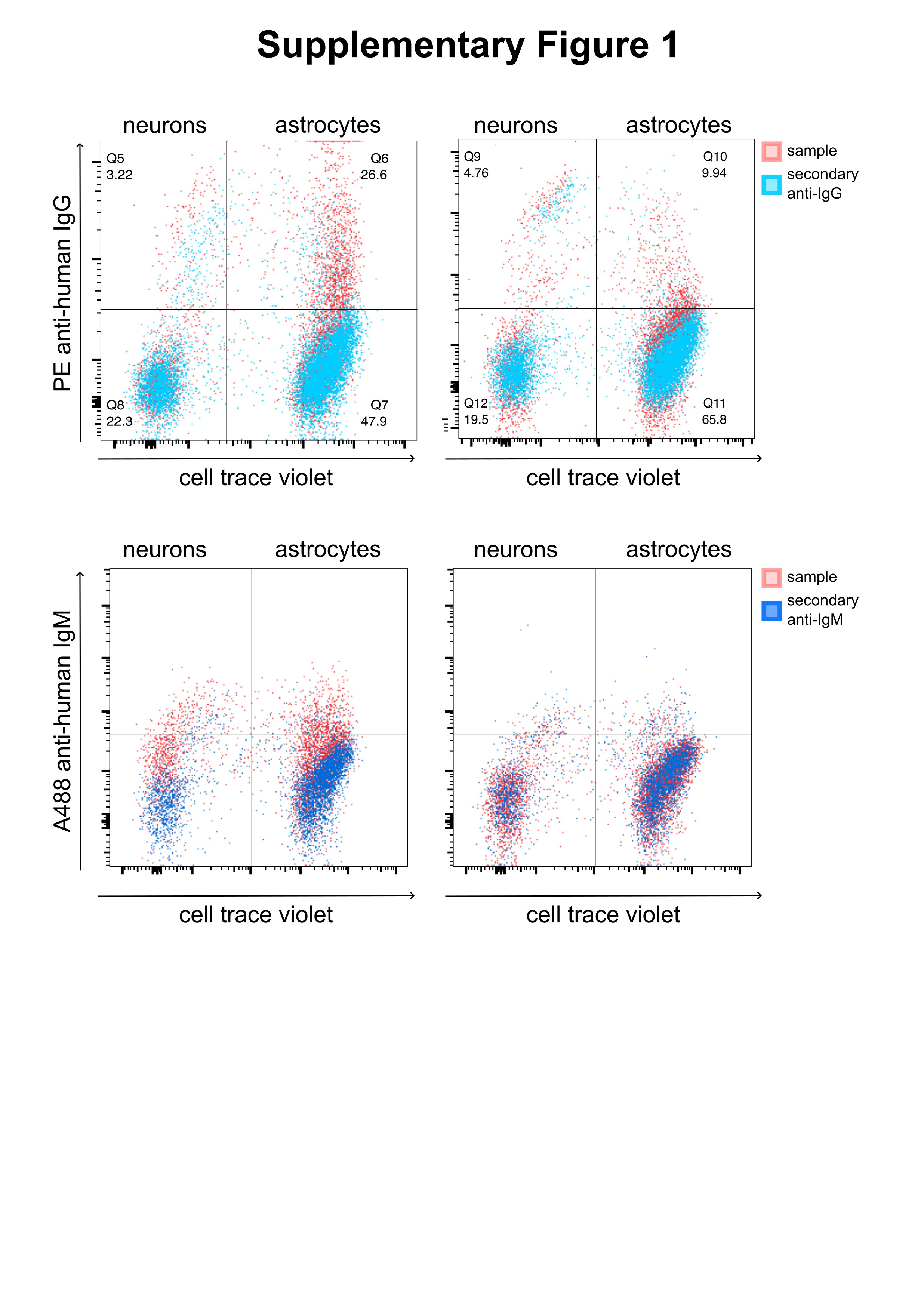

### Supplementary Figure 2

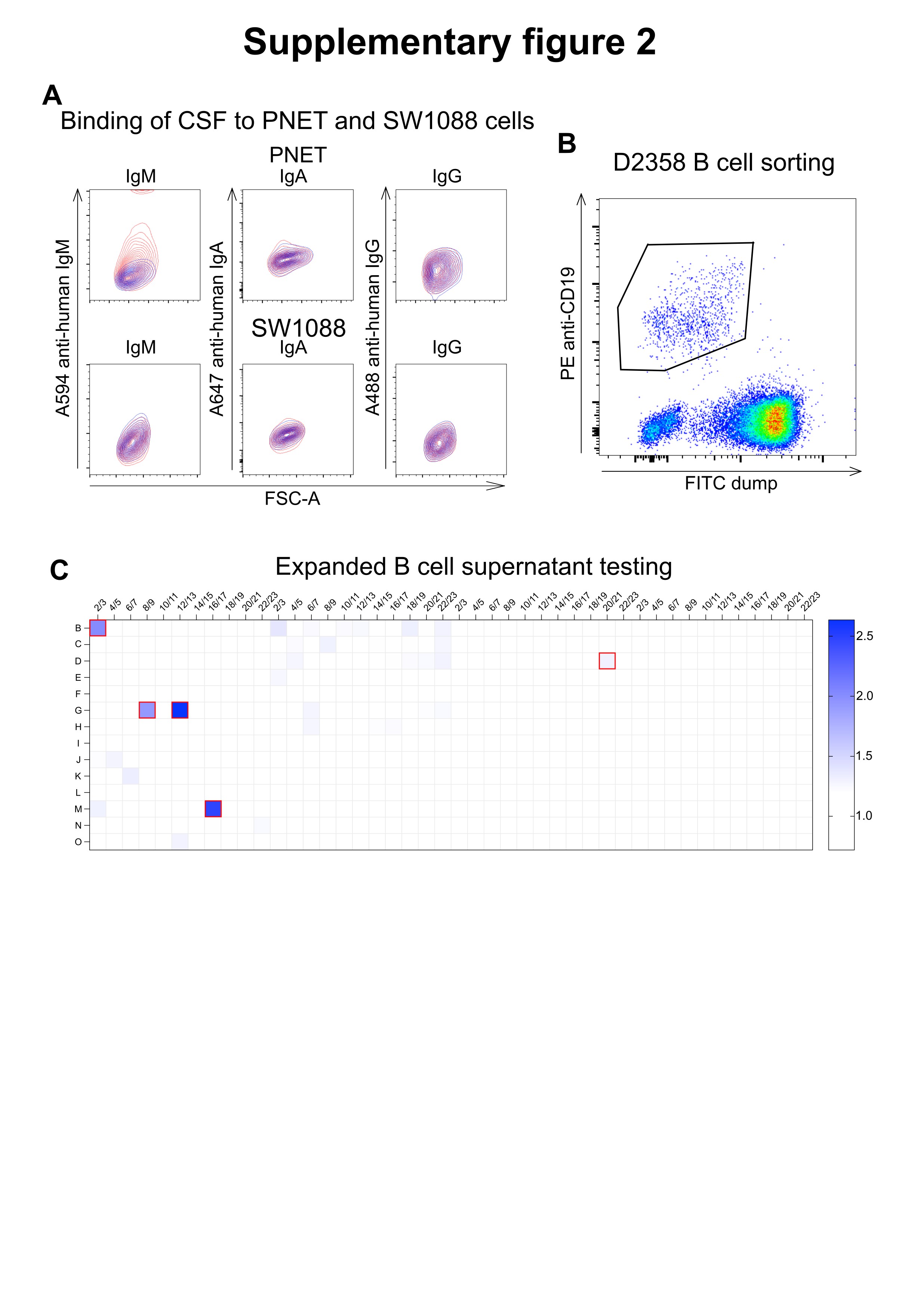

### Supplementary Figure 3

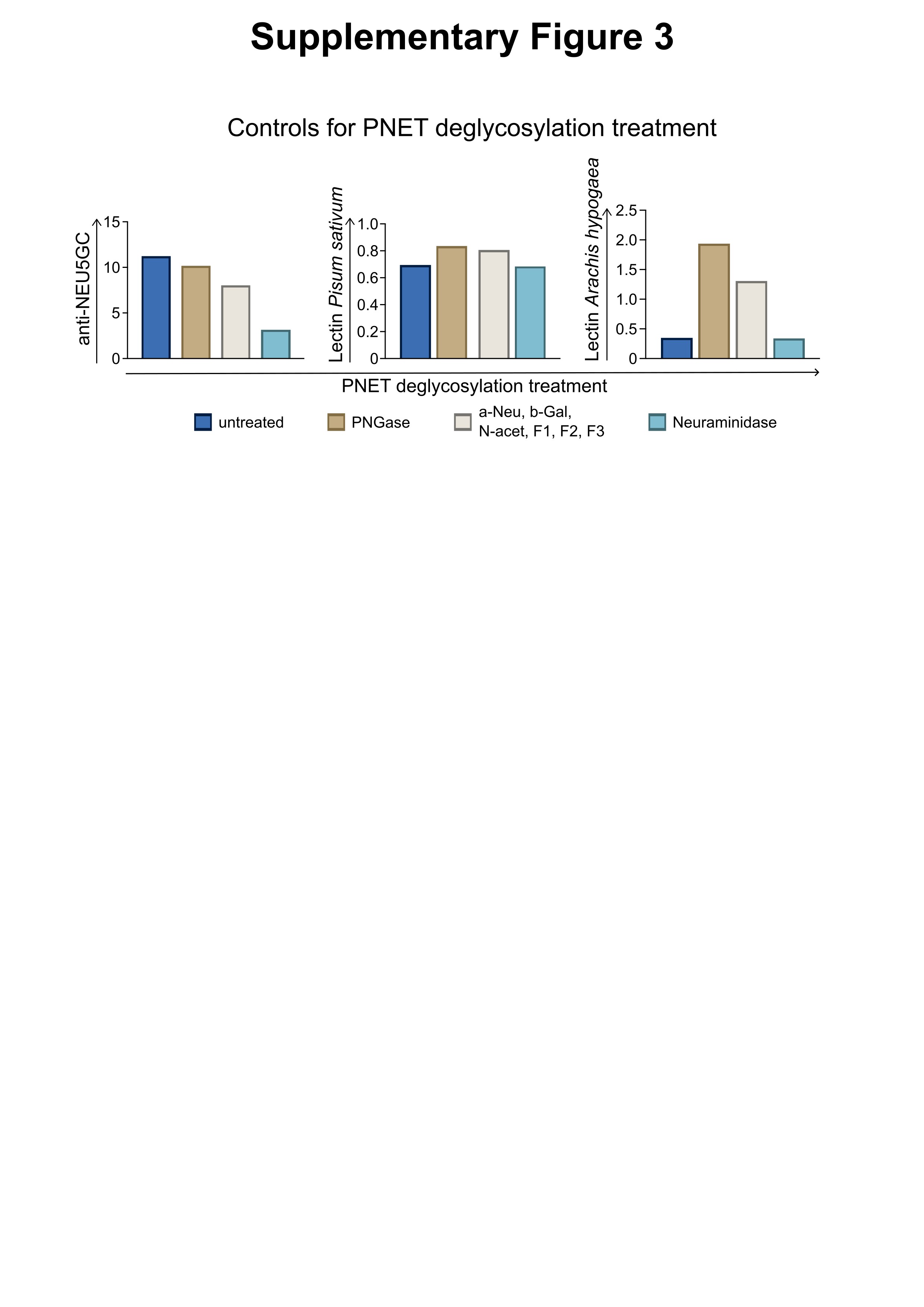

### Supplementary Figure 4

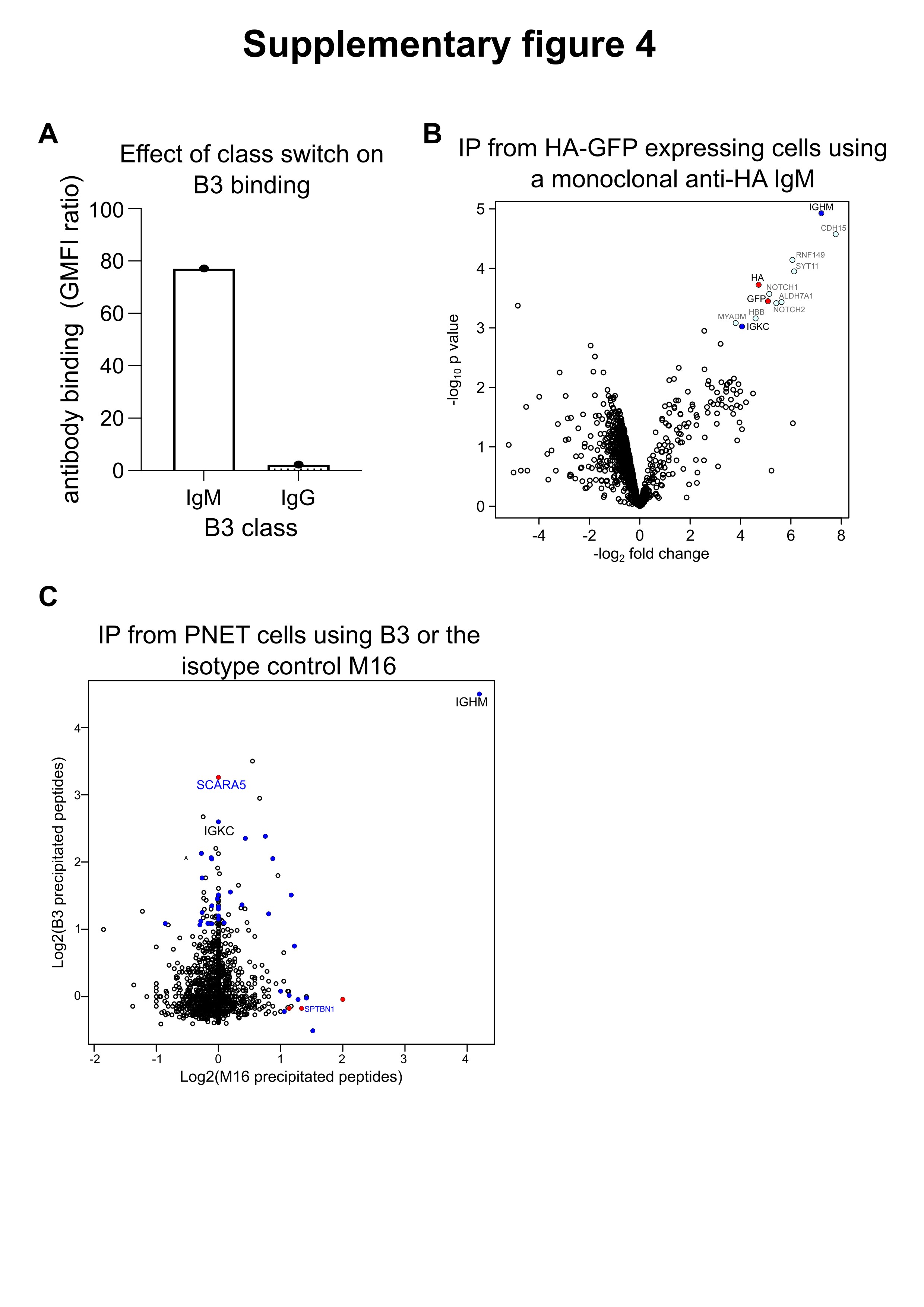

### Supplementary Figure 5

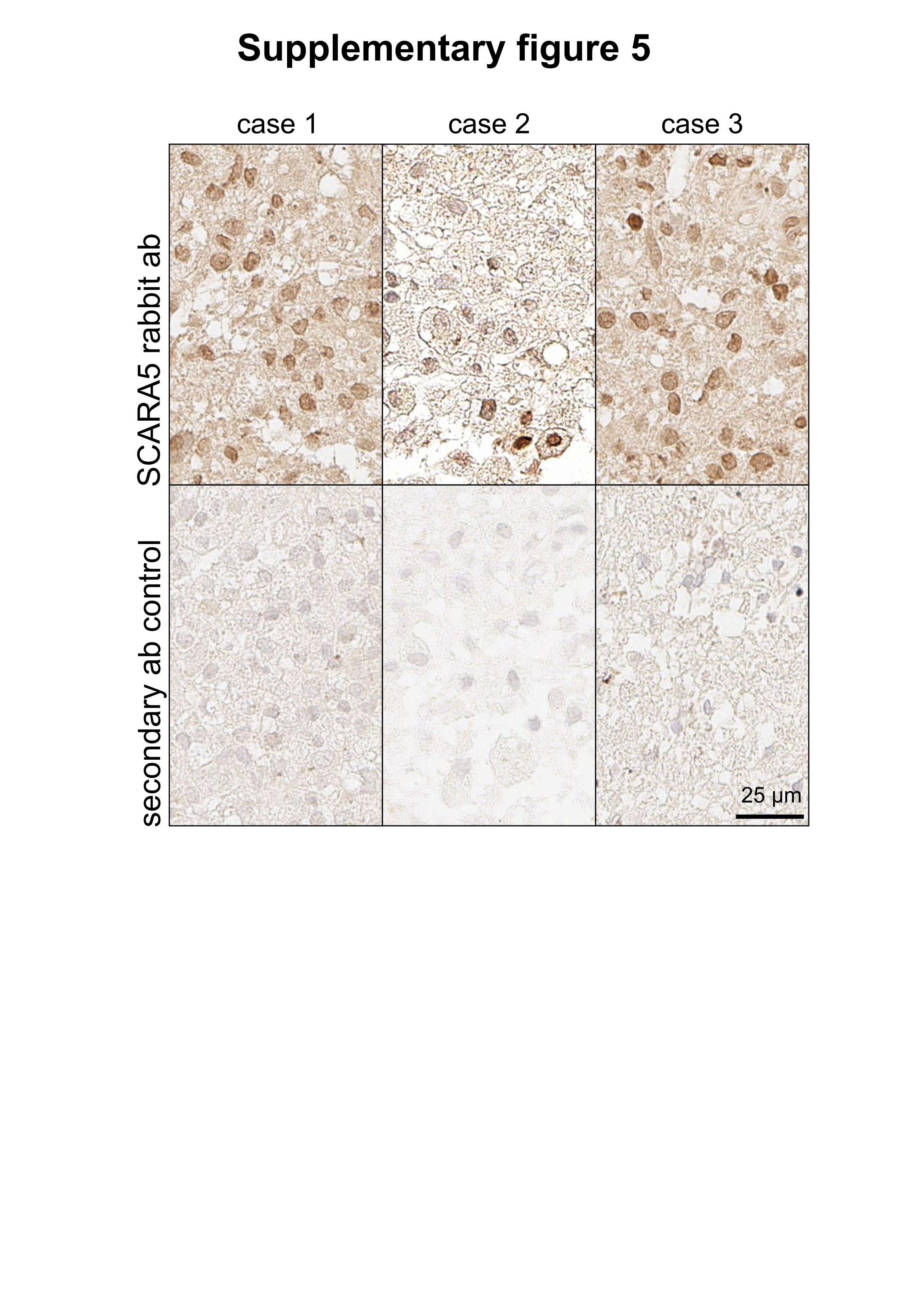
