## Supplementary Figure Ledends for "Cell-binding IgM in CSF is distinctive of multiple sclerosis and targets the iron transporter SCARA5"

### Supplementary Figure Legends

**Supplementary Figure 1**. CSF antibody binding to neurons (left side of each plot) or astrocytes (right side of each plot) derived from iPS cells. The vertical axis shows the fluorescence intensity of the anti-human IgG or IgM secondary, and the horizontal axis shows the intensity of the cell trace violet vital dye used to distinguish the two cell types. On each plot, the red dots show cells labeled with CSF followed by the anti-human IgG or IgM secondary antibody, and the blue dots cells labeled with secondary only. The plot on the top left shows results from a sample than contained astrocyte- binding IgG, and the plot on the right a CSF that contained no obvious cell-binding IgG. The plot on the bottom right shows results from a sample that contained both astrocyte- and neuron- binding IgM, and the plot on the right a CSF that contained no obvious cell-binding IgM.

**Supplementary Figure 2**. Isolation of PNET-binding monoclonal IgM antibodies from single CSF B cells. (**A**) Binding of three classes of antibody from the CSF of donor with PNET-specific IgM. Upper contour plots show binding to PNET, and lower plots binding to astrocytoma cell line SW1088. Red contours show results from CSF and blue contours results with secondary only. Vertical axis shows antibody binding (fluorescence intensity of secondary antibody) and horizontal axis shows forward scatter. (**B**) Gating strategy for single cell sorting. Cells from the CSF whose antibody binding profile is shown in A were labeled with anti-CD19 and a pool of FITC-labeled non-B cell markers, and the CD19-positive B cells sorted singly into wells. (**C**) Single B cells sorted as in (B) were incubated in the wells of a 384-well plate, in the presence of IL-21 and CD40L for 12 days, and then single well supernatants were tested for PNET binding in an analogous fashion to the CSF (A). Blue colour shown on bar at right represents the intensity of IgM PNET binding. Wells with significant PNET-binding IgM are highlighted with red squares, and B cells from these wells were used for the cloning procedure.

**Supplementary Figure 3**. Verification of efficiency and specificity of deglycosylation. The three bar charts depict assays of three different carbohydrate moieties: NEU5GC, detected with a commercial antibody; $\alpha$-mannose, detected with lectin from *Pisum sativum*; and β-gal(1→3)galNAc, detected with lectin from *Arachis hypogaea*. The colours of the bars indicate the enzymatic treatment, as shown beneath the figure.

**Supplementary Figure 4**. (**A**) Effect of artificial class switch on B3 binding to PNET cells. Either the native IgM (left bar) or an artificially class switched IgG version were used to label PNET cells, and binding quantified by flow cytometry. Vertical axis shows the GMFI ratio (antibody + secondary labeled cells: secondary only). (**B**) Immunoprecipitation and mass spectrometry to isolate protein target of known anti-haemagglutinin IgM. Cells transfected with haemagglutinin (influenza A/California/2009), fused to green fluorescent protein (GFP), were incubated with a recombinant monoclonal human-derived IgM (previously described by Zimmermann et al., 2019) against the extracellular domain of this membrane glycoprotein, and then bound proteins immunoprecipitated using magnetic beads coated with goat anti-human IgM antibodies. Precipitated proteins were digested with trypsin and subjected to mass spectrometry, and the numbers of peptides identified by this process plotted by gene. The horizontal axis shows the log2 fold difference between peptides precipitated by the anti-haemagglutin antibody and those from a control preparation in which the human anti-haemagglutinin IgM was omitted. The vertical axis shows the $\log_{10}$ p value from a Fisher's exact test, corrected for multiple comparisons by the method of Benjamini-Hochberg. Each black circle represents one protein. Red circles show the haemagglutin and GFP components of the precipitated antigen. Light blue circles are other, non-direct antigen proteins detected at a similar level, and dark blue circles show the heavy and light chains of the IgM. (**C**) Comparison of proteins immunoprecipitated by patient-derived IgM antibodies B3 and M16. Numbers of identified peptides were normalized for each antibody by dividing by the number of peptides identified in the same experiment under conditions of no IgM, and the $\log_2$ number of peptides identified from proteins immunoprecipitated by B3 was plotted against the same parameter for M16. Data from seven samples immunoprecipitated with B3 and six samples immunoprecipitated by M16 were averaged (arithmetic mean). Each point represents one protein. Membrane proteins are marked with solid blue circles. Those proteins from which significantly more peptides were immunoprecipitated by one antibody than by the other (Benjamini Hochberg False Discovery Rate < 0.05) are marked with solid red circles and their Gene Symbols have been added.

**Supplementary Figure 5. Localisation of SCARA5 in human brain.** Immunohistochemical staining of three additional early active MS lesion is shown. The top panels show labelling with SCARA5-antibody, the bottom panels show the secondary only control for each corresponding lesion.
