## Supplementary Table 1 for "Cell-binding IgM in CSF is distinctive of multiple sclerosis and targets the iron transporter SCARA5"

**Supplementary Table 1 Cell lines used**

| Name used in text | ATCC catalogue # | Cell type |
|---|---|---|
| STTG1 | CRL-1718 | Astrocytoma |
| SW1088 | HTB-12 | Astrocytoma |
| Daoy | HTB-186 | Medulloblastoma |
| A172 | CRL-1620 | Glioblastoma |
| PNET | CRL-2060 | neuroectodermal tumor |
| SVG p12 | CRL-8621 | fetal glial cells |
| HEK | CRL-11268 | embryonic kidney |
| TE671 | CCL-136 | rhabdomyosarcoma |
