## Supplementary Table 2 for "Cell-binding IgM in CSF is distinctive of multiple sclerosis and targets the iron transporter SCARA5"

**Supplementary Table 2 Demographic information about donors**

| | **Retrospective cohort (Basel)** | **Retrospective cohort (Graz)** |
|---|---|---|
| N of samples | 331 | 200 |
| CIS | 90 | 33 |
| MS | 78 | 67 |
| Controls | 163 | 100 |
| Age at sampling (y, median, range) | 47 (18-89) | 33 (18-70) |
| Sex, f (%) | 59% | 60% |
| Tested under immune treatment | 30 (9%) | 4 (2%) |
