## Supplementary Table 3 for "Cell-binding IgM in CSF is distinctive of multiple sclerosis and targets the iron transporter SCARA5"

**Supplementary Table 3 Clinical features of patients with CSF anti-PNET IgM**

| | **PNET binding IgM pos** | **PNET binding IgM neg** | |
|---|---|---|---|
| N | 36 | 291 | |
| MS subtype | | | |
| CIS | 19 | 126 | |
| RIS | 0 | 9 | |
| RRMS | 13 | 106 | |
| SPMS | 3 | 29 | |
| PPMS | 1 | 15 | |
| Age at sampling (y, mean, st dev) | **32 ± 12** | **42 ± 13** | **<0.0001** |
| Age at onset of first symptoms (y, mean, st dev) | **28 ± 12** | **36 ± 12** | **0.0003** |
| Sex, f (%) | 22 (61%) | 197 (68%) | 0.4280 |
| EDSS at sampling (median, interquartile range) | 2.0 (1.25) | 2.5 (2.0) | 0.3962 |
| Disease status, relapse (n, %) | 17 (47%) | 122 (41%) | 0.3682 |
| Immune treatment | | | |
| Untreated | 27 (75%) | 237 (82%) | 0.3551 |
| Injectable | 1 | 16 | |
| Oral | 3 | 13 | |
| Monoclonal | 5 | 17 | |
| Others | 0 | 8 | |
| CSF cell count n/uL (mean, st error) | **7 ± 1** | **5 ± 0.4** | **0.0269** |
| CSF albumin, mg/dl (mean, st error) | 215 ± 20.5 | 244 ± 10 | 0.2626 |
| CSF OCB | **36 (100%)** | **28 (79%)** | **0.0008** |
| PRL lesion count (median) | **2.5** | **2** | **0.5971** |
