## Supplementary Table 4 for "Cell-binding IgM in CSF is distinctive of multiple sclerosis and targets the iron transporter SCARA5"

**Supplementary Table 4 Immunoglobulin genes of antibodies cloned from CSF B cells**

| antibody | V genes (H & L) | J genes (H & L) | aa replacement (H & L) | specificity |
|---|---|---|---|---|
| B3 | V4-59 | J4*02 | 21 | PNET |
| B3 | KV2-28 | KJ1*01 | 4 | |
| D20 | V4-34 | J6*02 | 0 | nonspecific |
| D20 | LV3-19 | LJ3*02 | 0 | |
| G8 | V3-11 | J4*02 | 0 | nonspecific |
| G8 | LV3-25 | LJ2*01 | 0 | |
| M16 | V1-3 | J5*02 | 4 | nonspecific |
| M16 | LV3-23 | LJ1*01 | 3 | |
| G12 | V5-51 | J4*02 | 0 | nonspecific |
| G12 | LV3-1 | LJ1*01 | 0 | |
