## Supplementary Table 5 for "Cell-binding IgM in CSF is distinctive of multiple sclerosis and targets the iron transporter SCARA5"

**Supplementary Table 5 Genes overexpressed in B cell B3 compared to other CSF B cells**

| Gene Symbol | reads in B3 | average reads non B3 | ratio B3:average |
|---|---|---|---|
| GPR183 | 72 | 0.8 | 90 |
| KCNA3 | 34 | 0.4 | 85 |
| BHLHA15 | 23 | 0.4 | 57.5 |
| IRF4 | 80 | 1.6 | 50 |
| FCRL2 | 10 | 0.2 | 50 |
| CD74 | 901 | 19.8 | 45.5 |
| SASH3 | 18 | 0.4 | 45 |
| CD180 | 34 | 0.8 | 42.5 |
| TASL | 8 | 0.2 | 40 |
| HLA-DRB1 | 36 | 1 | 36 |
| LINC01854 | 6 | 0.2 | 30 |
| HLA-DQB1 | 28 | 0 | 29 |
| SYNE4 | 5 | 0.2 | 25 |
| KRT6B | 5 | 0.2 | 25 |
| CD79A | 89 | 3.6 | 24.7 |
| IGF1 | 9 | 0.4 | 22.5 |
| HLA-DRB5 | 20 | 0 | 21 |
| ZRANB2-AS1 | 4 | 0.2 | 20 |
| ATP13A4-AS1 | 4 | 0.2 | 20 |
| COL4A6 | 4 | 0.2 | 20 |
| IKZF3 | 24 | 1.2 | 20 |
| HLA-DRB3 | 4 | 0.2 | 20 |
| LY75 | 4 | 0.2 | 20 |
| ZDBF2 | 8 | 0.4 | 20 |
| RASSF6 | 26 | 1.4 | 18.5 |
| SELL | 10 | 0.6 | 16.6 |
| PLEK | 14 | 0 | 15 |
| SLFN11 | 14 | 0 | 15 |
| LINC02612 | 3 | 0.2 | 15 |
| SNX20 | 6 | 0.4 | 15 |
| ZNF296 | 6 | 0.4 | 15 |
| ZNF883 | 6 | 0.4 | 15 |
| DNASE2 | 24 | 1.6 | 15 |
| POU2AF1 | 27 | 1.8 | 15 |
| H2AJ | 6 | 0.4 | 15 |
| CA7 | 6 | 0.4 | 15 |
| C9orf64 | 6 | 0.4 | 15 |
| TNFSF8 | 3 | 0.2 | 15 |
| BMS1P20 | 157 | 10.8 | 14.5 |
| HLA-DRA | 13 | 0 | 14 |
| ARHGDIB | 32 | 2.4 | 13.3 |
| TNFRSF17 | 8 | 0.6 | 13.3 |
| IKZF1 | 10 | 0.8 | 12.5 |
| MAPK10 | 10 | 0.8 | 12.5 |
| PTPRCAP | 80 | 6.4 | 12.5 |
| PVRIG | 5 | 0.4 | 12.5 |
| CD53 | 60 | 4.8 | 12.5 |
| KRTAP21-2 | 11 | 0 | 12 |
| RHEX | 12 | 1 | 12 |
| RBM47 | 12 | 1 | 12 |
| LAX1 | 34 | 3 | 11.3 |
| KIAA0040 | 10 | 0 | 11 |
| MZB1 | 136 | 12.4 | 10.9 |
| ZBP1 | 19 | 1.8 | 10.5 |
