## Supplementary Table 6 for "Cell-binding IgM in CSF is distinctive of multiple sclerosis and targets the iron transporter SCARA5"

| host | catalog | isotype | immunogen | PNET | hSCARA5 | mSCARA5 | FFPE |
|---|---|---|---|---|---|---|---|
| mouse | MAB4900 | monoclonal IgG2b | NS0 mouse myeloma cell line transfected with human SCARA5 Arg83-His495 | + | + | ++ | NA |
| sheep | AF4754 | polyclonal IgG | Chinese hamster ovary cell line CHO-derived recombinant mouse SCARA5 Arg83-Pro491 | ++ | ++ | + | NA |
| rabbit | PA5-23551 | polyclonal IgG | KLH conjugated synthetic peptide between 385-413 amino acids from the C-terminal region of human SCAR5 | - | - | - | NA |
| rabbit | SAB3500071-100UG | polyclonal IgG | 17 amino acid peptide near the carboxy terminus of human SCARA5. | - | - | - | + |

**Supplementary Table 6.** Antibody binding to SCARA5 expressed on live cells, as assessed by flow cytometry, or on fixed tissue sections (FFPE) hSCARA5: Binding to HEK cells transfected with full-length (495 amino acid) human SCARA5. mSCARA5; Binding to cultured mouse brain cells. Specific binding estimated by comparing with no-primary controls. neither MAB4900, nor AF4754 compete for binding with B3 or with each other. Symbols: - no detectable binding; + detectable binding on some cells; ++ clearly delineated positive population; NA not tested.
